# Variation of tooth traits in ecologically specialized and sympatric morphs

**DOI:** 10.1101/2024.12.18.629189

**Authors:** Guðbjörg Ósk Jónsdóttir, Finnur Ingimarsson, Sigurður Sveinn Snorrason, Sarah Elizabeth Steele, Arnar Pálsson

## Abstract

Differences in dentition between species relate to feeding specialisations, as examples of tetrapod dentition variation show clearly. The association of tooth traits and ecological opportunties in non-mammalian vertebrates is less studied. We examined variation in dental traits in four sympatric morphs of Arctic charr (*Salvelinus alpinus*) which differ in feeding specialisations, head and jaw bone morphology. We studied tooth numbers in six bones (dentary, maxilla, premaxilla, palatine, vomer and glossohyal) and tooth angles in one bone (maxilla). We found fluctuating asymmetry in tooth numbers and angles and that the allometry of tooth numbers varied by bone but not morphs. Tooth numbers differed by morphs in four bones (dentary, palatine, vomer and glossohyal). In general, the morphs defined as pelagic had more teeth, and this relates partially to changes in bone shape. There was a difference in maxilla tooth angle, with benthic morphs having teeth which were angled more inwards. Dentary and maxilla tooth number correlated moderately with bone shape, maxilla tooth angle and premaxilla tooth number did not. While it is currently unknown what tooth characteristics are ancestral vs derived in these populations, the marked differences in specific bones presents an opportunity to explore rapid evolution in dentition.

**Statement and Declaration:** The authors declare that they have no conflict of interest

## Introduction

Adaptive evolution in vertebrates can entail substantial degrees of changes and variations on feeding behaviour and how organisms process food (Bels et al., 1994; Gidmark et al., 2019; Schwenk, 2000). Phenotypic adaptations in feeding elements have been linked to vast diversification in many vertebrate groups via prey specialization (Gidmark et al., 2019; Grant & Grant, 2003). Dentition is an example of this as variation in teeth traits associate with various feeding specialisations in vertebrates (Gregory et al., 2016; Huysseune, 1995; Koussoulakou et al., 2009; Melstrom, 2017; Streelman et al., 2003). Many aspects of dentition, for example shape and inclination angle, can affect their efficiency in fulfilling their main roles (catch, hold or process prey, Anderson & LaBarbera, 2008; Crofts & Summers, 2014). Therefore, many aspects of tooth traits vary, like number (Koussoulakou et al., 2009; Line, 2003), complexity (Melstrom, 2017), shape (Gregory et al., 2016; Streelman et al., 2003) and location in the jaw (Mihalitsis & Bellwood, 2019) and this is often related to prey capture and processing. Among the best examples of ecologically specialized dentition are illustrated in extant and fossil mammals (Line, 2003), but similar adaptations and specialisations have also been found in non-mammalian vertebrates, like reptiles (Lafuma et al., 2021; Melstrom, 2017) and fishes (Carr & Motta, 2020; Huysseune, 1995; Streelman et al., 2003).

Most fish are homodont (i.e. similar shaped teeth, Berkovitz & Shellis, 2023a; Kolmann et al., 2019) and constantly replace their teeth (polyphyodonty, Carr et al., 2021; Fraser et al., 2006; Huysseune & Witten, 2008; Kolmann et al., 2019). This differs from mammals which are generally heterodont (i.e. differently shaped teeth (Ungar, 2010) and replace their teeth only once throughout their life (diphyodonts). While majority of fish are classified homodont variation in their dentation is known (Berkovitz & Shellis, 2023a; Mihalitsis & Bellwood, 2019). For example, tooth numbers can vary with size (Cleves et al., 2014; Ellis et al., 2015; Greer, 1991; Torres-Carvajal, 2007) or between left and right on paired bones, showing asymmetry (Witten et al., 2005). The number of teeth generally increase allometrically as the individual grows (Cleves et al., 2014; Ellis et al., 2015; Greer, 1991; Torres-Carvajal, 2007). However, allometric relationships can differ depending on species and the bone carrying the teeth (Greer, 1991; Torres-Carvajal, 2007), with these relationships also possibly being plastic (Huysseune, 1995). While bilateral symmetry is the general pattern in vertebrates (Klingenberg et al., 2002; Koeberle et al., 2020) individual organisms are rarely perfectly symmetric (Klingenberg et al., 2002). For tooth numbers the degree of asymmetry can vary by species, bone (P. Currie, 2003; P. J. Currie, 2003; Miyashita & Currie, 2010) and even throughout the individual’s the life cycle (Witten et al., 2005). Asymmetry is generally defined as: 1) directional - where the left and right sides differ in a systematic way (Palmer & Strobeck, 1986; Van Valen, 1962), 2) antisymmetry – bimodal distribution of left and right differences about a mean of zero (Palmer & Strobeck, 1986; Van Valen, 1962), and 3) fluctuating - non-directional deviations from perfect symmetry (Merilä & Björklund, 1995; Palmer & Strobeck, 1986; Van Valen, 1962). Directional asymmetry is thought to have a genetic basis and be developmentally controlled (Palmer & Strobeck, 1992; Van Valen, 1962). However, fluctuating asymmetry stems from developmental noise generated by chance events, manifest in interactions of the genotype and environmental variation throughout ontogeny (Koeberle et al., 2020; Merilä & Björklund, 1995; Van Valen, 1962).

Fishes show great diversity in form and function of dentition, with many diverse shapes, like conical teeth which can pierce prey, molariforms used for grinding and crunching, and broad blade-like teeth useful for slicing (Berkovitz & Shellis, 2023a; Gidmark et al., 2019). Variation in tooth number, both inter- and intraspecific, is also rather common in fish (Cleves et al., 2014; Eastman, 1977; Ellis et al., 2015; Huysseune, 1995; Märss et al., 2017), with differences in number often being linked to diet (Huysseune, 1995). Intraspecific differences in tooth number have been shown in many fish lineages, for example in cichlidae (Huysseune, 1995) and three-spined sticklebacks (*Gasterosteus aculeatus*, Caldecutt et al., 2001; Cleves et al., 2014; Ellis et al., 2015). Rearing experiments indicate that tooth numbers have both a heritable component (Cleves et al., 2014), and can also be plastic (Huysseune, 1995; Karagic et al., 2020; Worcester, 2012), with different numbers of teeth being associated with different diet treatments (Huysseune, 1995). For example, in the cichlid species *Astatoreochromis alluaudi*, where snail feed fish have fewer teeth on the pharyngeal jaw than fish feed on softer prey (Huysseune, 1995).

The functional importance of dentition is clear as teeth increase the efficiency of the jaw in capturing prey and often process it as well (Berkovitz & Shellis, 2023b). Variation in many aspects of dentition, be it shape (Crofts & Summers, 2014), inclination angle (Anderson & LaBarbera, 2008), number (Huysseune, 1995) or location in the jaw (Mihalitsis & Bellwood, 2019), could be adaptations to different prey items (Crofts & Summers, 2014; Huysseune, 1995; Mihalitsis & Bellwood, 2019). For example, inclination angle can affect cutting efficiency (Anderson & LaBarbera, 2008) or efficiency in holding prey (Dean et al., 2008; Gosline, 1973) and an increase in tooth number in three-spined sticklebacks has been linked to possible adaptations for crushing larger prey (Cleves et al., 2014). However, dentition may also diverge neutrally due to genetic drift, or as correlated response to other adaptations via linkage or pleiotropy (Cleves et al., 2014). Teeth count generally appear to be linked with the size of the dental palate (tooth-bearing section), tooth spacing (Cleves et al., 2014; Ellis et al., 2015) and tooth shape (Huysseune & Witten, 2018). They may also associate with less directly related traits, through genetic correlations. A QTL (quantitative trait loci) affecting tooth spacing in sticklebacks overlaps with the Ectodysplasin (Eda) gene (Cleves et al., 2014) which controls adaptive reductions in armour plates (Colosimo et al., 2005). Therefore, changes in dental and other traits could associate with tooth number.

After the last glacial retreat, many new freshwater habitats opened in the Northern Hemisphere. Many waters were colonized by anadromous fish species, with some populations staying anadromous (traveling between freshwater and the sea) while others became landlocked. These freshwater habitats where usually species poor (minor interspecific competition) but had a diversity of unutilized habitats and food resources (Snorrason & Skúlason, 2004) and resource polymorphisms often arose in these system (Smith & Skúlason, 1996; Snorrason & Skúlason, 2004). Therefore, these systems provide an excellent opportunity to study intraspecific phenotypic variations and possible adaptive divergence (Snorrason & Skúlason, 2004). In this paper we study one such salmonid system, the four Arctic charr (*Salvelinus alpinus*) morphs in lake Þingvallavatn, in Iceland.

Salmonids are known for extensive phenotypic variability and in the case of Arctic charr polymorphism and plasticity (Klemetsen, 2010; Noakes, 2008). Arctic charr shows extensive phenotypic variation throughout its distribution (Klemetsen, 2010), and sympatric morphs that differ in feeding ecology are found in many lakes (Doenz et al., 2019; Skoglund et al., 2015; Woods et al., 2013; Østbye et al., 2020). Within lake Þingvallavatn four Arctic charr morphs coexist (Malmquist et al., 1985; Sandlund et al., 1992). Two of the morphs primarily feed on benthic invertebrates (large benthivorous (LB) and small benthivorous (SB) charr) and two morphs capture prey in the water column (planktivorous (PL) and piscivorous (PI) charr), here after called benthic and pelagic morphs for simplification (Malmquist, 1992). These morphs differ in many life-history and ecological traits, for example habitat use, adult size and spawning times (Jonsson et al., 1988; Malmquist, 1992; Malmquist et al., 1985; Sandlund et al., 1987; Skúlason, Noakes, et al., 1989; Skúlason, Snorrason, et al., 1989; Snorrason et al., 1989). The morphs notably differ in feeding ecology, with LB- and SB-charr feeding mainly on snails (mainly *Radix balthica*), PL-charr on crustacean zooplankton and PI-charr mostly on three-spined sticklebacks (Malmquist, 1992). Differences in feeding ecology may have influenced divergence in spawning times among these morphs (Brachmann et al., 2021; Skúlason, Snorrason, et al., 1989; Smith & Skúlason, 1996). The differences in feeding ecology have also been linked to the substantial variation in head morphology (Snorrason et al., 1989). Our recent study also found differences in the shape of four oral jaw bones which appears linked to feeding ecology (Jónsdóttir et al., 2024). Three of these four bones are tooth-bearing jaw bones (Day, 1887). Therefore, as differences in morph prey preference appear to be linked to morphological variation, we wanted to study tooth trait divergence.

Rather few studies have explored tooth trait variation in salmonids, though tooth development and replacement has been described in rainbow trout (*Oncorhychus mykiss*, Fraser et al., 2006) and Atlantic salmon (*Salmo salar* L., Huysseune & Witten, 2008). Variation in teeth of Arctic charr has received some attention, with reports of differences between morphs in glossohyal tooth number during development (Pichugin, 2009) and population differences in tooth number and length on some bones in adults (Bryce et al., 2016). Within the lake Þingvallavatn morphs, an honours thesis revealed several patterns, i) positive allometric relationship of fish size and teeth numbers (on maxilla, palatine, vomer and glossohyal), ii) differences in tooth number among morphs in the dentary and palatine (pelagic morphs having more teeth) and iii) while maxilla tooth numbers did not vary by morph, the inward inclination (measured as an angle) of the teeth was significantly greater in benthic morphs (Ingimarsson, 2002). We revisit tooth variation in the specialized Þingvallavatn morphs with a larger sample size to investigate the variation and divergence of tooth traits. We expect increased numbers of teeth in pelagic morphs which feed on more evasive prey and expect the highest number of teeth within the PI-charr which feeds on the larges prey (Cleves et al., 2014). Maxilla teeth angle is expected to differ between morphs, with benthic fishes having more inward inclination allowing them to hold their prey while sediment is cleaned (Dean et al., 2008). We examined the tooth number and angle in the Arctic charr morphs of Þingvallavatn. As variation in dentition may relate to ecological differences by morphs in prey capture and handling, we test if variation in these tooth traits differs by bones and relates to variation in body size, ecological specialization, and shape of tooth-bearing bones among morphs, all known to relate to feeding ecology. We ask:

1. Do tooth traits (numbers and inclination angle) show fluctuating asymmetry (FA), and if so, does FA vary with fish size or morph?
2. Is there an allometric relationship between tooth traits and body size, and does this differ by bones and morphs?
3. Do tooth numbers per bone and angles of the maxillary teeth differ by morph?
4. Does tooth number and inclination angle correlate with the axes of shape of the respective and other bones?

## Methodology

### Tooth Imaging and Counts

Two datasets were utilized. The primary analyses were conducted on a large dataset of Arctic charr (N=240) caught in 2020 and 2021, consisting of adults of all four morphs from Lake Þingvallavatn (see details of sampling and processing in (Jónsdóttir et al., 2024). To examine if patterns were consistent over time, data from a study mentioned prior, (N=138) on the same morphs collected in 1997 (Ingimarsson, 2002) was examined, here after referred to as the “second dataset”. All tooth counts and related data are provided as supplements (Online Resources 1 to 42) and as datafiles with analyses pipeline on Github (https://github.com/GudbjorgOskJ/CharrTeethTTV).

For the primary dataset, oral jaw bones (dentary, maxilla, premaxilla, palatine, vomer and glossohyal) were photographed (Canon EOS 77D, 50 mm Microlens) in stereotypical view with scale and teeth analysed. The dentary and maxilla were photographed in medial and ventral view, premaxilla in fronto-lateral view, palatine in lateral and medial view, vomer in lateral and ventral view and glossohyal in lateral and dorsal view (Fig. 1). Teeth were counted twice on each bone: March and May 2023, by same researcher (GOJ). The focus was on fully ankylosed teeth, however as some teeth fell out during processing, we also counted broken teeth and holes left from broken teeth as teeth (Fig. 1b). For the symmetric bones (dentary, maxilla, premaxilla and palatine) we had four replicate measures (left-right, March and May) and two for the vomer and glossohyal. Counting was highly repeatable (Online Resources 2 - 4). We used available landmark data for dentary, maxilla and premaxilla bones from Jónsdóttir et al. (2024) to examine relationships between tooth traits and bone shape. All pictures of fish and bones are available on Figshare+ (doi:10.25452/figshare.plus.25118825).

**Fig. 1.**
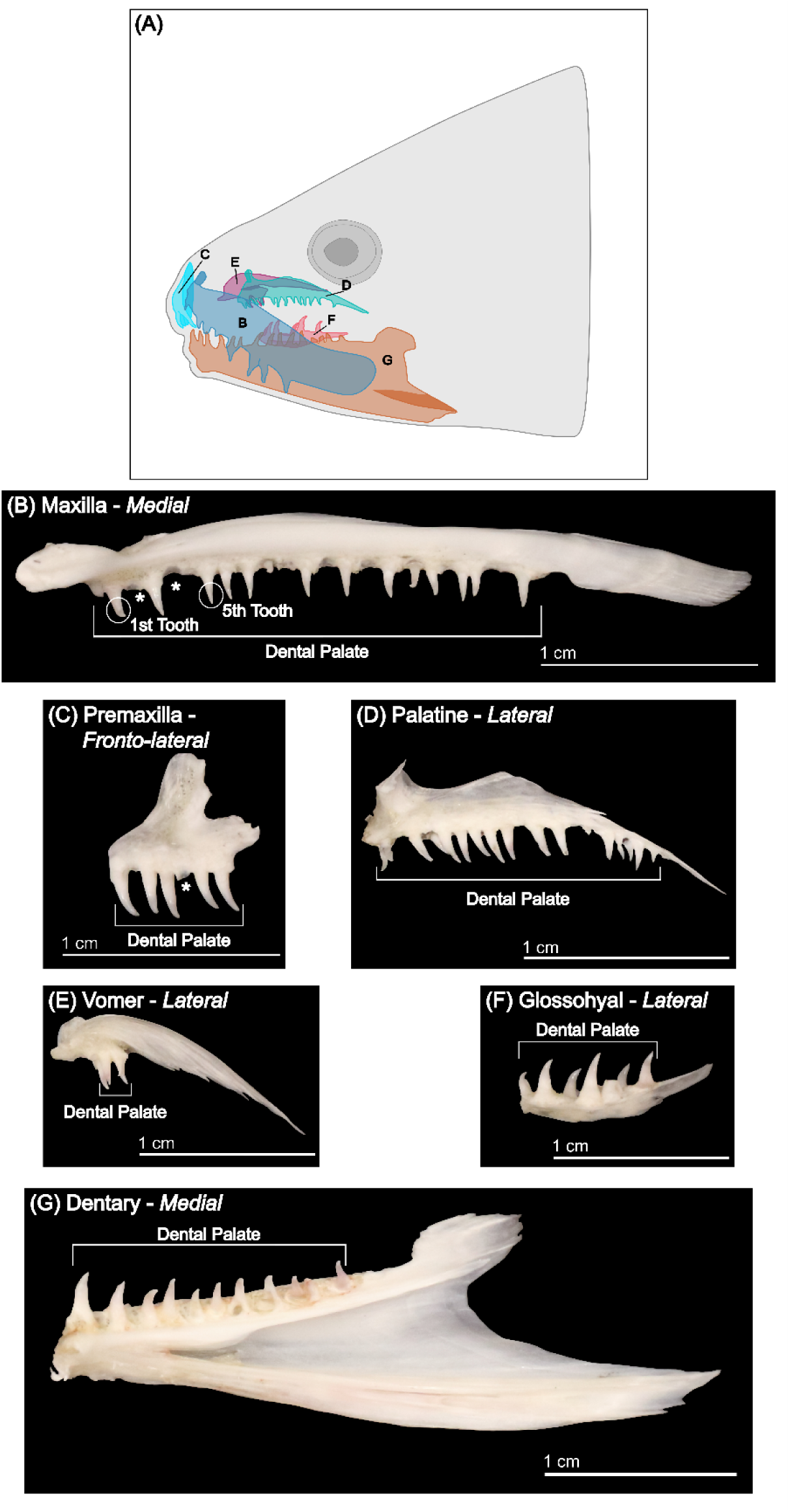
Morphology of the six craniofacial bones in Arctic charr studied here. (A) Locations of the six bones drawn on a schematic of a charr head, with front facing left. Note this is schematic figure, bones are not too scale. B-G) Images of the jaw bones used, with names and angle of photographing (italic). Dental palate (defined as distance from first tooth to last tooth) is indicated on each bone. Shown are bones of individual PC4386 (PI-charr, L = 36.6 cm and W = 435 g, male), inset scale (1 cm). B-C) White stars show examples of missing teeth, that were counted as true teeth here (may have fallen out during processing). Approach for measuring maxilla tooth angles is demonstrated in Online Resource 1.

For the “second dataset”, jaw bones (dentary, maxilla, premaxilla, palatine, vomer and glossohyal) were originally photographed (JVG camcorder, model E 5506DC), with scale in stereotypical view from both sides (left and right jaws). The dentary was photographed in medial view, maxilla in ventral view, premaxilla in fronto-lateral view, palatine in lateral view, vomer in lateral and glossohyal in dorsal view. Teeth were counted once for one side by the same researcher (FI) at the time of the initial study (Ingimarsson, 2002). We only had access to the flat file with recordings (deposited on Github, https://github.com/GudbjorgOskJ/CharrTeethTTV), as the raw image files were not available.

### Maxilla tooth angles

To calculate the angles of maxillary teeth of the primary dataset, we photographed the maxilla again using a stereomicroscope in three views: medial, ventral and fronto-lateral (focus on 1^st^ tooth, Online Resource 1). To develop this trait, we tested two methods to measure angles. First, we used two photographs (medial and ventral view, here called “medial/ventral” method). The lengths of two teeth (1^st^ and 5^th^) were measured from both medial and ventral view. These teeth were chosen for convenience and comparison to “fronto-lateral” method (see below). From these two measurements the tooth angle was calculated using the tangent:

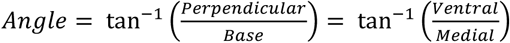

Second, we used a “fronto-lateral” method developed by Ingimarsson (2002). Using fronto-lateral view photographs of the maxilla we assessed the angle (1^st^ tooth) directly with linear measures in Fiji (Schindelin et al., 2012). Repeatability of both methods was tested on a random subsample of ∼60 individuals of three morphs (LB-, PI- and PL-charr, ∼20 for each). Repeatability of the fronto-lateral method and medial/ventral method was tested on a subsample, 62 individuals, with three of the morphs (LB-, PI- and PL-charr, ∼20 for each morph). Linear regression showed that the medial/ventral method was more repeatable (Online Resources 23 and 24). Therefore, tooth angle data was collected with the medial/ventral method. All measurements were done by the same researcher. The “second dataset” came from photographs of the maxilla bones (again JVG camcorder, measured by FI at the time of the initial study, (Ingimarsson, 2002). The angle was measured directly with Fiji on the tooth which was in best focus (the 2^nd^ to 5^th^ tooth) on each bone.

### Statistical analyses of tooth counts

All statistical analyses were done in R (vs. 4.2.3, R Core Team, 2023) in R-studio (vs. 2023.06.1+524, RStudio, 2023). Figures were generated in R and formatted with Inkscape (vs. 1.3 (0e150ed6c4, 2023-07-21), Inkscape, 2020). Body size of individuals was estimated with fork length (FL, ln transformed). The repeatability of counts was assessed with a linear regression model (function *lm*), for all bones, left and right side independently (when applicable) and the mean (average number of teeth per fish). Pearson correlation was also estimated (*cor.test*).

To examine asymmetry in tooth numbers in the paired bones, we ran *glm* models on the differences between the left and right bone. The asymmetry in tooth numbers was calculated per individual (left – right, for the symmetric bones), which also allows estimation of the proportions of individuals with “perfect” symmetry (same number of teeth on both sides). Then, we asked whether these deviations correlated across bones (using *cor*), and if they deviated from 0 (directional asymmetry), varied by morphs, changed with body size (FL) and if asymmetry-allometry varied by morphs, with linear models in R:

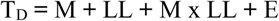

TD: deviation in count, M: morph, LL: ln(FL). We used Bonferroni correction for four bones, α = 0.01.

The variation in tooth numbers on the six bones was analysed using a generalized linear mixed model (GLMM) implemented with the Template Model Builder (TMB) (the *glmmTMB* package in R, version 1.1.7, Brooks et al., 2024). First, we tested the impact of morph, sex, and estimates of size (ln(FL)) on all bones together and then each separately, assuming a gaussian distributed response variable (tooth number). The models were as follow:

All bones model:

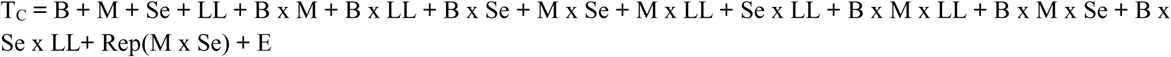

By bone model:

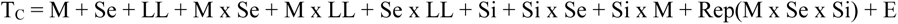

In these models, TC: tooth count, B: bone, M: morph, Se: sex, LL: ln(FL), Si: side (left or right bone), with some interaction terms. Note, we accounted for repeated measures as a random term nested under fixed effects. Also, for bones with Centroid size estimates (C-size for dentary, maxilla and premaxilla, from geometric morphometric analyses, see Jónsdóttir et al. (2024)) we also ran the model with ln transformed C-size. As we studied six bones, a Bonferroni corrected threshold α = 0.05/6 = 0.008 was used. For the “second dataset” we ran the same model (dropping the side and replicate counts variables) to test effects of morph, sex and size.

### Statistical analyses of maxilla tooth angles

All individuals in the primary dataset were scored with the medial/ventral method and all in the “second dataset” with the “fronto-lateral” method (differences discussed below). To examine the influence of morph, sex and size on maxilla tooth angles we again ran a generalized linear mixed model (GLMM) (*glmmTMB* package) with a gaussian distribution. The model was similar as above, except tooth was a random variable (1^st^ and 5^th^), but not repeated per individual. Again, we used two proxies for size, ln FL, and ln C-size of the maxilla. As angle was only analysed on maxilla, we used α = 0.05. For the “second dataset” we ran the same model (dropping the side and tooth variables).

### Tests of shape and tooth trait covariation

We assessed the covariation of tooth traits (counts and angles) and shapes of tooth bearing bones (Jónsdóttir et al., 2024). Using two-block partial least squares (PLS) with the R function *two.b.pls* (*geomorph*), we compared tooth traits and shape of the same and other bones, both for all morphs together and each separately. Effect sizes were compared between bones (and between morphs within an individual bone), with *compare.pls.* For this analysis we used α = 0.007 as a conservative threshold (Bonferroni correction for 7 traits, 6 tooth counts and 1 angle measurement).

## Results

### Fluctuating asymmetry and variation in tooth numbers and allometry by bones

We counted tooth numbers on the dentary, maxilla, premaxilla, palatine, vomer and glossohyal, and estimated the angle of inward inclination of the maxilla teeth (see below). There appeared to be some fluctuating asymmetry in tooth numbers in the four paired bones (Fig. 2 and Online Resource 5), with a minority of individuals having the same number of teeth on both sides (from 15% for dentary to 21% for premaxilla). However, there was no indication of directional asymmetry, as left minus right tooth number did not differ from zero (Online Resource 8). Neither was their indication of directional asymmetry by size, morph nor their interaction (Online Resources 9 and 10). Finally, there was no correlation of these deviations across bones (Online Resource 11).

**Fig. 2.**
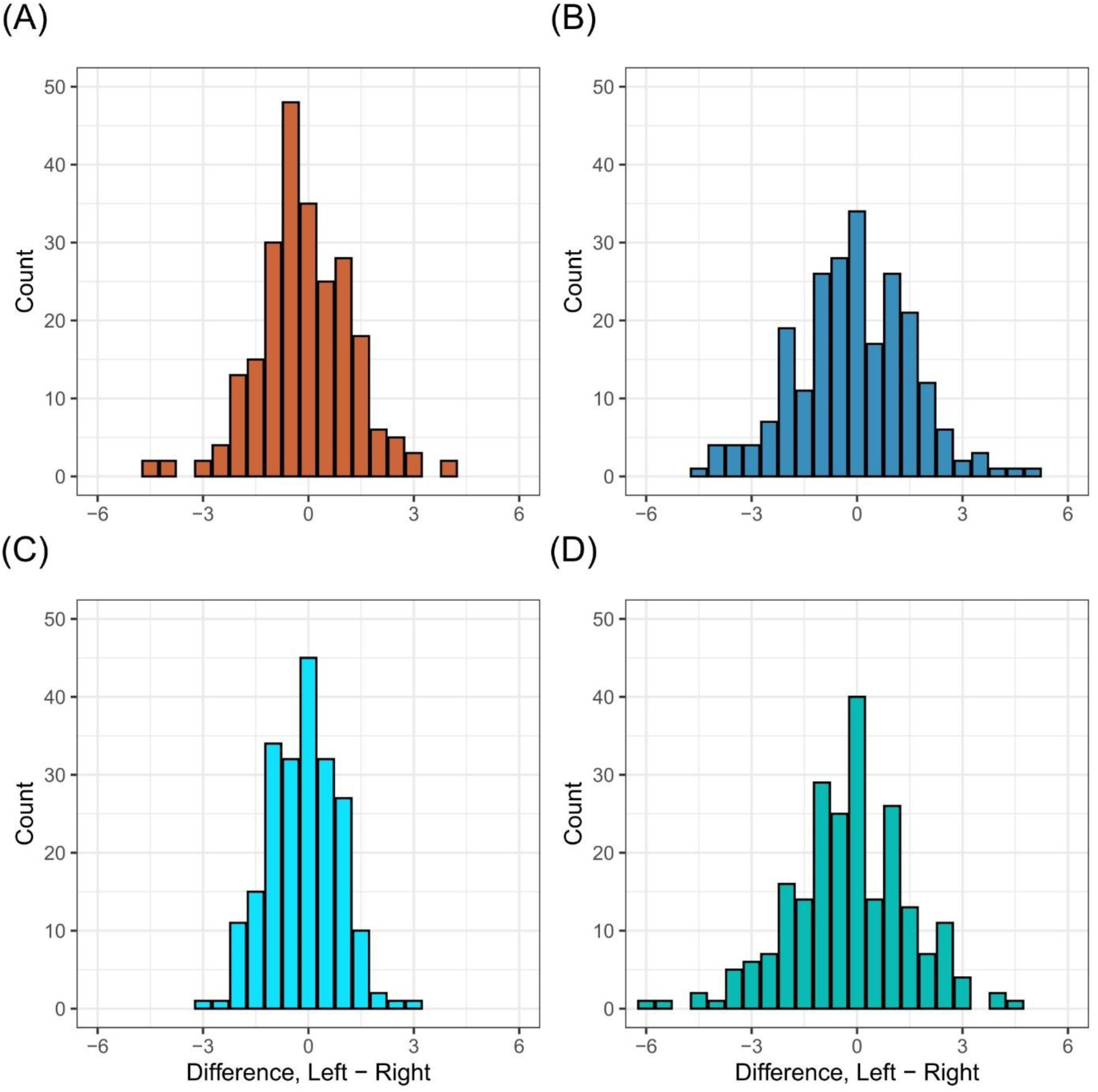
Fluctuating asymmetry in tooth numbers on four paired Arctic charr bones. Shown are histograms of the within individual variation, as the deviation in tooth numbers between the sides (subtracting right from left), for the dentary (A), maxilla (B), premaxilla (C) and palatine (D). X-axis shows the deviation between left and right and the Y-axis the number of individuals per bin. Note the scale is continuous, as averages from two measures per individual are represented.

As expected, the numbers of teeth varied greatly by bone (Fig. 3). The average tooth number, over all morphs and both sexes, was 14.7 for the dentary (range 9 to 26), 20.0 (12 to 28) for the maxilla, 7.5 (4 to 10) for the premaxilla, 16.5 (9 to 24) for the palatine, 4.5 (1 to 9) for the vomer and 12.0 (8 to 20) for the glossohyal (Online Resource 12). This reflects differences in types and sizes of bones, or possibly the dental palate (tooth-bearing part) - like in the dentary (Fig. 1g). When analysed by morphs, the dentary had the widest range, from average 12.79 teeth in SB-charr to 16.96 in PI-charr. The narrowest range was in the premaxilla (7.26 in LB-charr to 7.61 in PI-charr). Predictably, bones with many teeth also had the highest variance (standard deviation for maxilla was 2.26, compared to the premaxilla at 0.7 teeth). To study these apparent morph differences, we fitted mixed models (that utilized the repeated measures) with size as a covariate (cause the morphs differ greatly in size, (Sandlund et al., 1992).

**Fig. 3.**
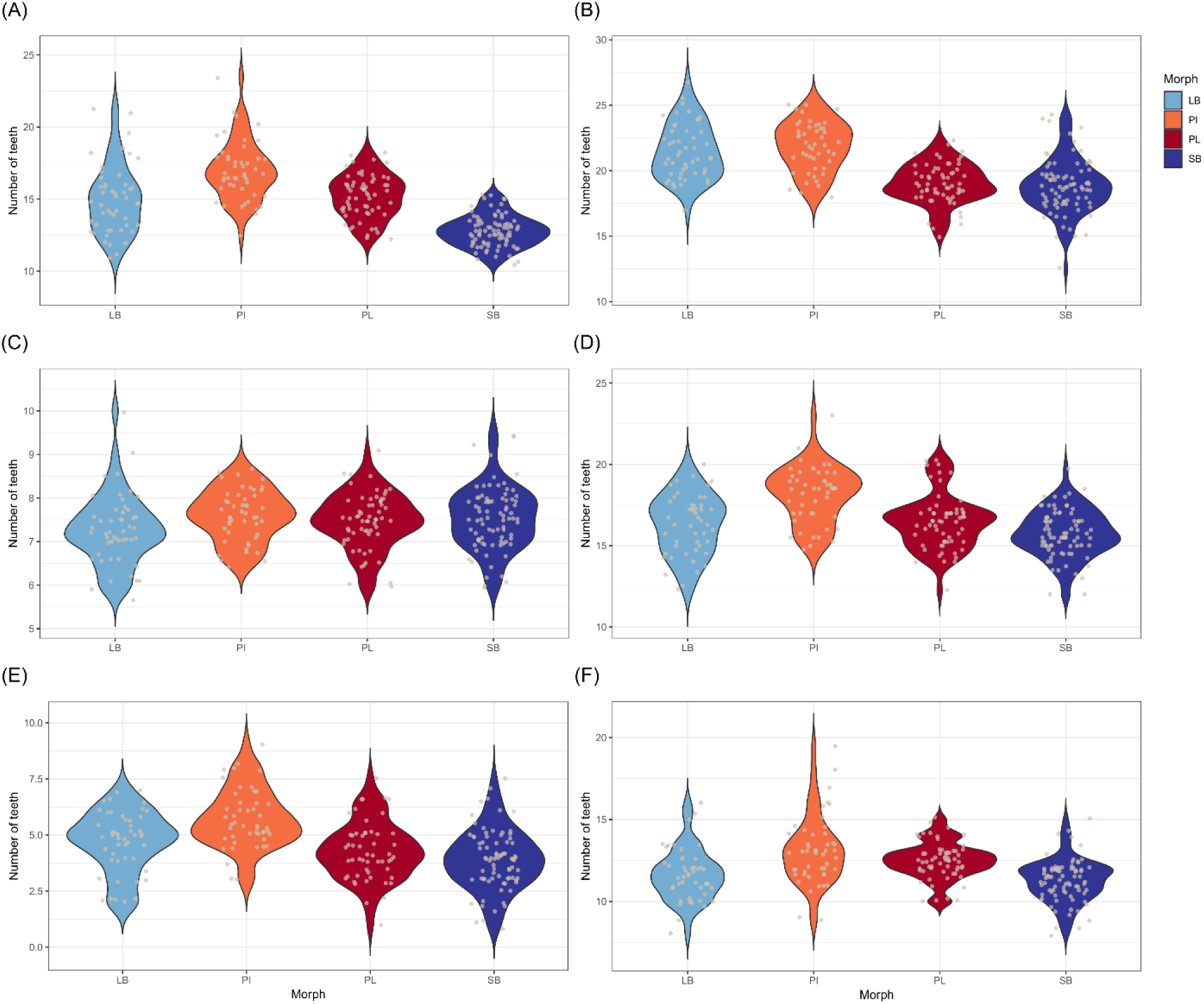
Distribution of tooth number in six bones by morph. The Y-axis indicates the number of teeth (average per individual for the paired bones A-D, average over repeated measures for the non-paired E-F). The bones are: dentary (A), maxilla (B), premaxilla (C), palatine (D), vomer (E) and glossohyal bone (F). Morphs are coloured: SB: dark blue, LB: light blue, PI: orange, PL: red (see also inset).

We explored the variation in tooth numbers using both a mixed model framework and an ANOVA on size corrected tooth numbers (see methods). First, tooth numbers varied by bone, and we found significant interaction of bone with morph and with size (Table 1). There were some examples of three-way interactions also. This confirms the at least partial independence of tooth numbers on distinct bones, which led us to conduct bone by bone analyses.

**Table 1.** The influence of bone, size, morph, sex and side (left or right) on tooth number in six jaw bones. Analysed with generalized linear mixed models (glmmTMB) using ln FL for size. P-values below α = 0.005 are shown in bold.

| Variable | $\chi^2$ | Df | p-value |
| --- | --- | --- | --- |
| Bone | 56506.799 | 5 | < **0.001** |
| Morph | 61.121 | 3 | < **0.001** |
| Sex | 0.040 | 1 | 0.842 |
| Size (ln FL) | 32.350 | 1 | < **0.001** |
| Bone x Morph | 369.425 | 15 | < **0.001** |
| Bone x Sex | 9.958 | 5 | 0.076 |
| Bone x ln FL | 64.692 | 5 | < **0.001** |
| Morph x Sex | 4.383 | 3 | 0.223 |
| Morph x ln FL | 3.227 | 3 | 0.358 |
| Sex x ln FL | 0.171 | 1 | 0.680 |
| Bone x Morph x Sex | 36.518 | 15 | **0.001** |
| Bone x Morph x ln FL | 50.461 | 15 | < **0.001** |
| Bone x Sex x ln FL | 9.737 | 5 | 0.083 |

### Size and morph affect tooth numbers on most bones but not all

For most bones, the mixed models revealed strong and consistent effects size and morph on tooth numbers, but sex only had a minor effect, and no two-way or higher order interaction terms were significant (Table 2). Body size strongly influenced tooth numbers in all bones, except premaxilla and glossohyal (Table 2, Fig. 4). The slope was steepest for the dentary and palatine (Online Resource 15), as an increase of an individual’s length by 5 cm led to ∼0.5 and ∼0.4 extra teeth respectively. Analyses with centroid size gave the same results (for the three bones with such estimates, (Jónsdóttir et al., 2024), Online Resource 16). The analyses did not detect interaction of size with morph. Similarly, there did not appear to be consistent sexual dimorphism for tooth numbers (Table 2, sex effects minimal to non-existent) and no sex interaction term was significant (Table 2). The only trait with sex dependence was tooth angle (see below). More interestingly, the models indicated clear morph differences in four bones (dentary, palatine, vomer and glossohyal), largest for the dentary. In general, tooth numbers were highest in the large PI-charr and fewest in the small SB-charr (on average over all bones, Fig. 3 and Online Resource 13). The complementary analyses, ANOVA on size corrected data, also showed morph differences. The LB-charr had the fewest teeth in most bones, and SB-charr had the highest numbers (Online Resource 14). The post hoc tests indicated stronger differences across all bones between the larger morphs (LB- < PI-charr), than the smaller ones (SB- and PL-charr). Finally, the “second dataset” (fish collected in 1997) gave generally consistent results, with the pelagic morphs having more teeth on average (Online Resources 34 - 37). These patterns will be described for each bone.

**Fig. 4.**
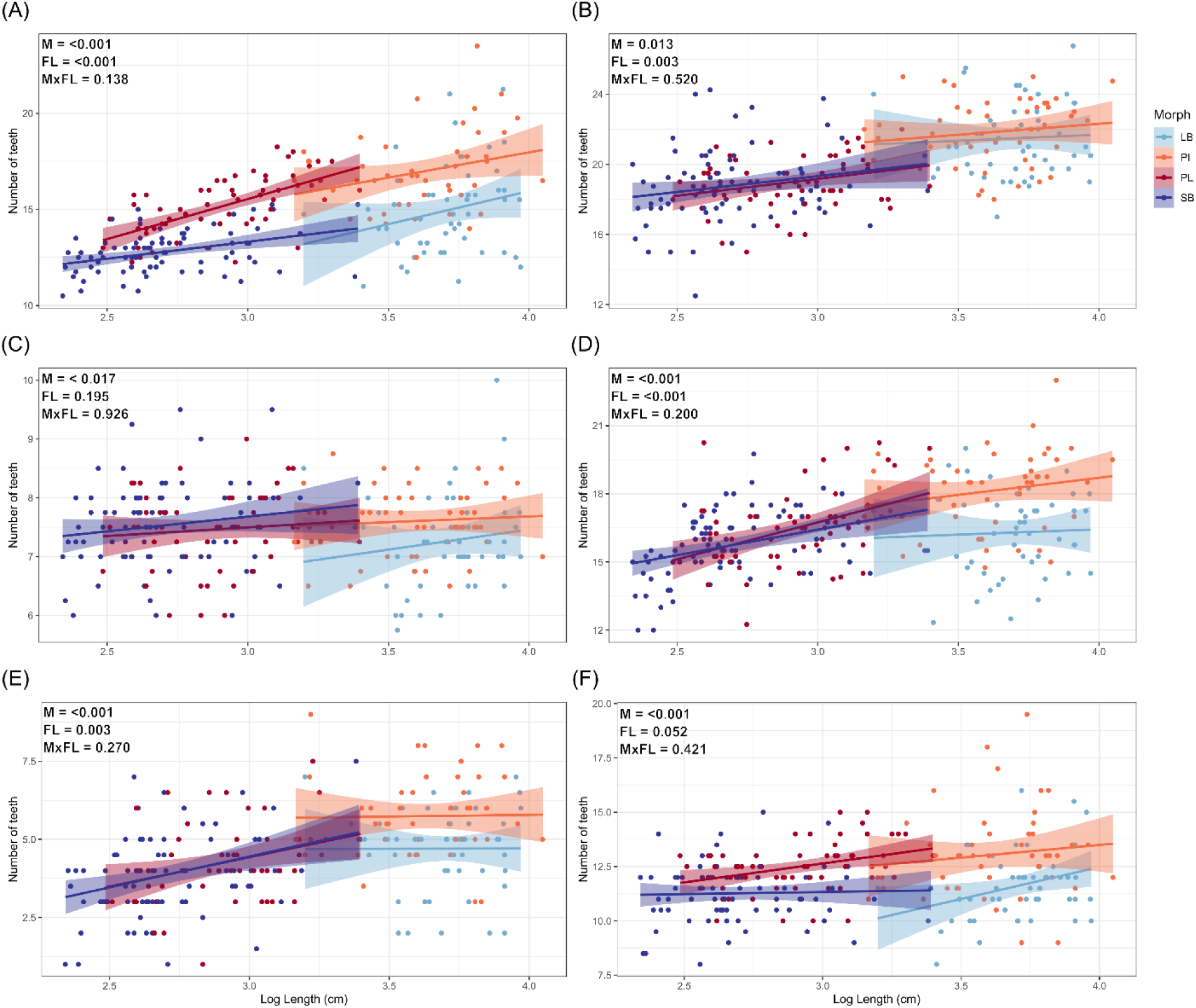
Influence of size and morph on number of teeth varied by bone. The scatterplots show individual tooth numbers (Y-axis) against size (ln(FL in cm) X-axis) for the four morphs and six bones; A) dentary, B) maxilla, C) premaxilla, D) palatine, E) vomer and F) glossohyal. Note the numbers for symmetric bones (A-D) are averages for the left and right bones. Each plot shows p-values from glmmTMB for the effect of morph (M), body size (FL) and morph by body size (MxFL) on tooth number.

**Table 2.** The influence of size, morph, sex and side (left or right) on tooth number in six jaw bones. Analysed with generalized linear mixed models (glmmTMB) using ln FL for size. P-values below the Bonferroni corrected threshold (α = 0.008) are shown in bold

| Terms |  | Dentary | Maxilla | Palatine | Premaxilla | Vomer | Glossohyal |
| --- | --- | --- | --- | --- | --- | --- | --- |
| Morph | $\chi^2$ | 91.072 | 10.700 | 44.715 | 10.153 | 19.508 | 46.465 |
|  | df | 3 | 3 | 3 | 3 | 3 | 3 |
|  | p-value | **< 0.001** | 0.013 | **< 0.001** | 0.017 | **< 0.001** | **< 0.001** |
| Sex | $\chi^2$ | 0.406 | 0.121 | 0.030 | 4.907 | 2.090 | 0.036 |
|  | df | 1 | 1 | 1 | 1 | 1 | 1 |
|  | p-value | 0.524 | 0.728 | 0.862 | 0.027 | 0.148 | 0.851 |
| FL (ln FL) | $\chi^2$ | 31.124 | 8.747 | 18.434 | 1.677 | 8.814 | 3.769 |
|  | df | 1 | 1 | 1 | 1 | 1 | 1 |
|  | p-value | **< 0.001** | **0.003** | **< 0.001** | 0.195 | **0.003** | 0.052 |
| Side | $\chi^2$ | 1.745 | 0.910 | 4.909 | 5.273 | NA | NA |
|  | df | 1 | 1 | 1 | 1 | NA | NA |
|  | p-value | 0.187 | 0.340 | 0.027 | 0.022 | NA | NA |
| Morph x Sex | $\chi^2$ | 2.899 | 6.019 | 1.783 | 4.144 | 1.485 | 2.517 |
|  | df | 3 | 3 | 3 | 3 | 3 | 3 |
|  | p-value | 0.407 | 0.111 | 0.619 | 0.246 | 0.686 | 0.472 |
| Morph x ln FL | $\chi^2$ | 5.519 | 2.261 | 4.638 | 0.289 | 3.920 | 2.816 |
|  | df | 3 | 3 | 3 | 3 | 3 | 3 |
|  | p-value | 0.138 | 0.520 | 0.200 | 0.962 | 0.270 | 0.421 |
| Sex x ln FL | $\chi^2$ | 0.075 | 1.072 | 0.008 | 0.642 | 0.000 | 1.483 |
|  | df | 1 | 1 | 1 | 1 | 1 | 1 |
|  | p-value | 0.785 | 0.301 | 0.928 | 0.42305 | 0.992 | 0.223 |

NA: Each fish only has one copy of these bones, hence no “Side” term.

The dentary is the anterior-most part of the lower jaw and its tooth numbers were affected both by size and morph (p < 0.001), but the interaction term was not significant. The benthic fish had fewer teeth than pelagic fish matching in size. The pairwise tests (over both sexes) indicated significant differences between PL- and SB-charr (p < 0.001) and between LB- and PL-charr (p < 0.001), with PL-charr having ∼3 more teeth than both SB- and LB-charr. The difference between LB- and PI-charr was ∼2.3 teeth but non-significant (p = 0.02, Online Resource 17). Palatine and vomer showed similar patterns, with significant size and morph effects on tooth numbers (more teeth on average in pelagic fish, Table 2). However, for neither bone did the pairwise tests of morphs reveal differences (Online Resources 17 and 18), suggesting subtler differences than in the dentary. The glossohyal tooth numbers were strongly affected by morph, but not significantly by size. Again, the pelagic morphs had higher tooth numbers, shown by pairwise tests (significant differences between LB- and PL-charr, p = 0.003, with PL-charr having ∼2.5 more teeth). The LB-PI-charr comparison pointed in the same direction (PI-charr > LB-charr) but was non-significant (unadjusted, p = 0.012, Online Resource 17). For both the maxilla and premaxilla (in upper jaw like the palatine), morph did not significantly affect tooth numbers. Maxilla tooth numbers associated positively with size, however curiously, premaxilla tooth numbers did not.

### Morph differences in maxilla tooth angles

We explored the influence of morph, sex and size on maxillary tooth angles with *glmmTMB*. The strongest effects appeared to be morph, sex and an interaction of morph by tooth (difference between 1^st^ and 5^th^ tooth). First, the inward inclination of maxillary teeth differed by morph, with the mean angle (over both teeth) being largest in SB-charr (50.9°) but smallest in PL-charr (32.4°). Differences in this trait among morphs was along a benthic-pelagic axis, as the benthic morphs had more inward angled maxillary teeth. Pairwise tests showed significant differences between the LB- and PI-charr, with LB-charr having more inward angled teeth (p = 0.01, Online Resource 26) as well as between the LB- and PL-charr (p < 0.001) and SB- and PL-charr (p < 0.001) always with the benthic morphs having more inward angled teeth (Online Resource 26). Second, while size influenced maxilla tooth numbers, it did not relate to their angles (Table 3, Online Resource 29). Third, several other terms were significant for maxilla tooth angle, notably the asymmetry, the two teeth measured and sex. Side (left vs right, estimate of asymmetry) was significant, with the left side being slightly more angled inward on average (estimated 2° differences, p < 0.001), but no interaction with side was significant (Table 3). Tooth angle did not differ between the 1^st^ and 5^th^ tooth, as the average angle, over the entire dataset, was 43.8° for both the 1^st^ and 5^th^ maxilla tooth. However, there was significant interaction between the tooth and morph effects (p = 0.001). For the larger morphs (LB- and PI-charr) the 5^th^ tooth was angled more inward, but for the smaller morphs (PL- and SB-charr) the 1^st^ tooth was angled more inward. Within morphs this difference was only significant in the PI-charr (estimated 2.7° differences, p = 0.017) and PL-charr (estimated 2.7° differences, p = 0.01). Males tended to have less angled teeth (p < 0.001), this sex difference appeared largest in the PL-charr, morph by sex interaction is non-significant (p > 0.07, Fig. 5 and Online Resources 25, 27 and 28). This was the only tooth trait indicating sex differences.

**Fig. 5.**
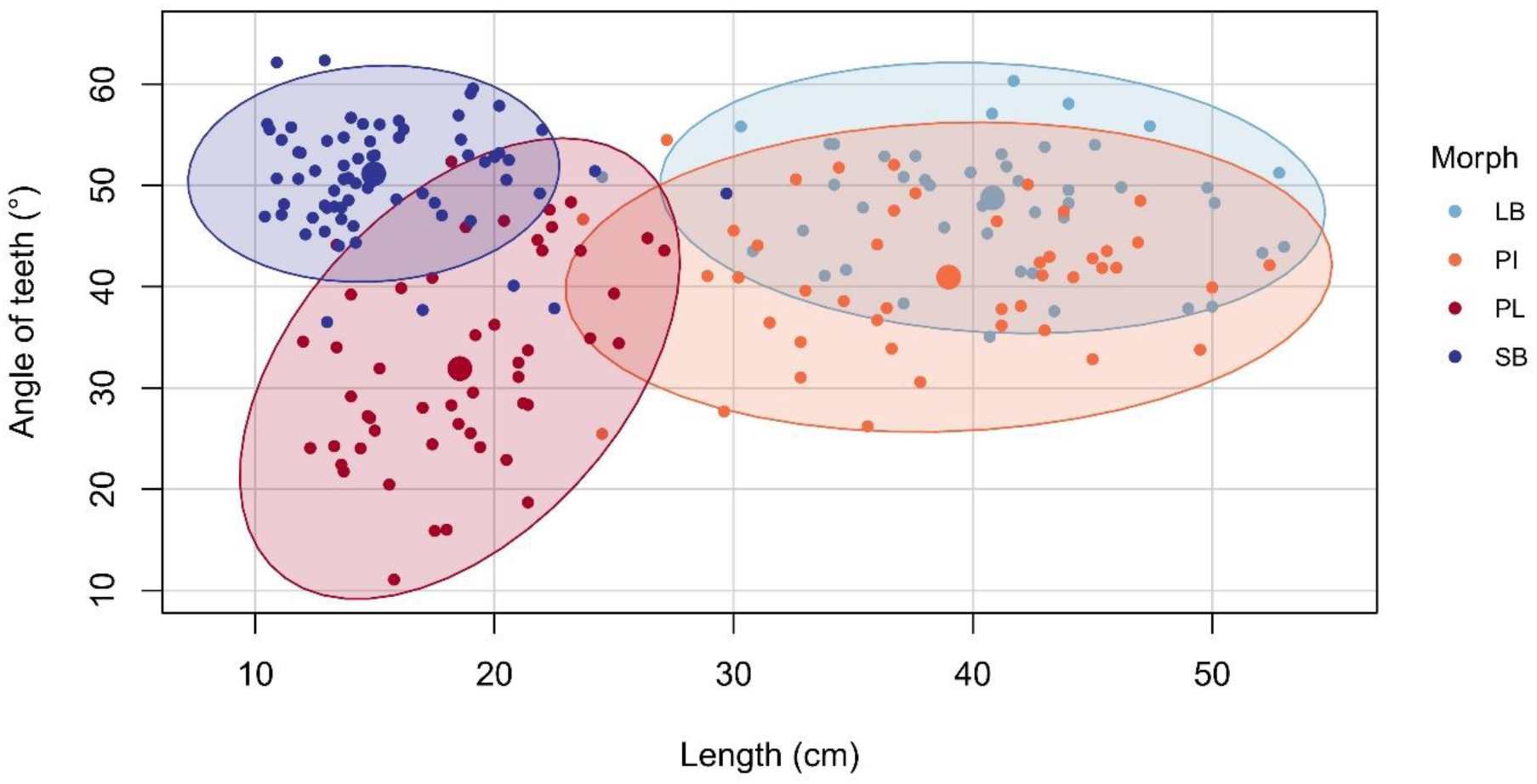
Effect of morph on the inward inclination of maxillary teeth in the four morphs. The scatterplot shows average tooth angle (for 1^st^ and 5^th^ tooth on both left and right bone, Y-axis) against size (fork length of fish, X-axis) for all individuals. The ellipses and shaded areas are 95% CI for the distributions and large dots represent means per morph.

**Table 3.** The influence of size, morph, sex, side (left or right) and tooth (1^st^ or 5^th^ tooth) on the inward inclination of maxillary teeth. Analysed with generalized linear mixed models (glmmTMB) using ln (FL) for size

| Variable | $\chi^2$ | df | p-value |
| --- | --- | --- | --- |
| Morph | 252.101 | 3 | **< 0.001** |
| Sex | 25.919 | 1 | **< 0.001** |
| Size (ln FL) | 0.222 | 1 | 0.638 |
| Side | 16.375 | 1 | **< 0.001** |
| Tooth (1<sup>st</sup> v. 5<sup>th</sup>) | 0.001 | 1 | 0.978 |
| Morph x Sex | 7.071 | 3 | 0.070 |
| Morph x ln FL | 3.439 | 3 | 0.329 |
| Sex x ln FL | 1.360 | 1 | 0.244 |
| Morph x Tooth | 16.416 | 3 | **0.001** |

Analyses of the “second dataset” showed the same patterns, the benthic morphs having more inward angled maxillary teeth than PL-charr (Online Resources 38 - 40). The mean angles in the groups differed substantially, between the datasets. Most likely this reflects methodological differences as the maxillary angles were measured with two different methods in the two datasets.

### Patterns of covariation between tooth traits and bone shape

Lastly, we explored the covariation of tooth traits (counts and angles) on particular bones and the shape of the same bone (and others as a reference). We calculated the correlation coefficient from two block partial least squares, r-PLS, for the entire dataset and morphs separately. For the entire dataset, nearly all pairs of bones and tooth traits were significant, with the degree of covariation generally being low to moderate (r-PLS ∼ 0.38 – 0.72, p < 0.001, Online Resource 41). The strongest association was between dentary shape and dentary tooth number (r-PLS = 0.717, p < 0.001). The lowest values were the premaxilla tooth numbers, that did not covary with premaxilla, dentary or maxilla shape (Online Resource 41), likely reflecting low variation in this trait. We concluded the pattern of covariation in the overall dataset reflect differences in bone size, shape and/or tooth numbers between morphs, and thus analysed the data by morph. Predictably, we found a lot fewer significant pairs of bone shape and bone tooth numbers. These covariations were primarily between tooth numbers and shape of the corresponding bone. For instance, dentary shape and dentary tooth number showed moderate but significant level of covariation in the two smaller morphs (SB- and PL-charr, r-PLS = 0.52 and 0.69 respectively, p = 0.001, Fig. 6 and Online Resource 42) and the maxilla shape covaried significantly and moderately with maxilla tooth number in two morphs (LB- and PL-charr, r-PLS = 0.53 and 0.55 respectively, p ≤ 0.004, Fig. 6 and Online Resource 42). Again, premaxilla shape did not covary with premaxilla tooth number (in any morph). Other relationships were generally weak and non-significant, with some exceptions. Maxilla shape and dentary tooth numbers correlated in PI- and PL-charr (r-PLS = 0.54 and 0.62 respectively, p ≤ 0.005, Table 4), dentary shape and palatine tooth numbers in PL-charr, and premaxilla shape and maxilla tooth numbers for SB-charr (r-PLS = 0.529, p = 0.003) and glossohyal tooth numbers for PI-charr (r-PLS = 0.599, p = 0.001). Finally, maxilla tooth angles did not covary with shape of any bone (Online Resource 42).

**Fig. 6.**
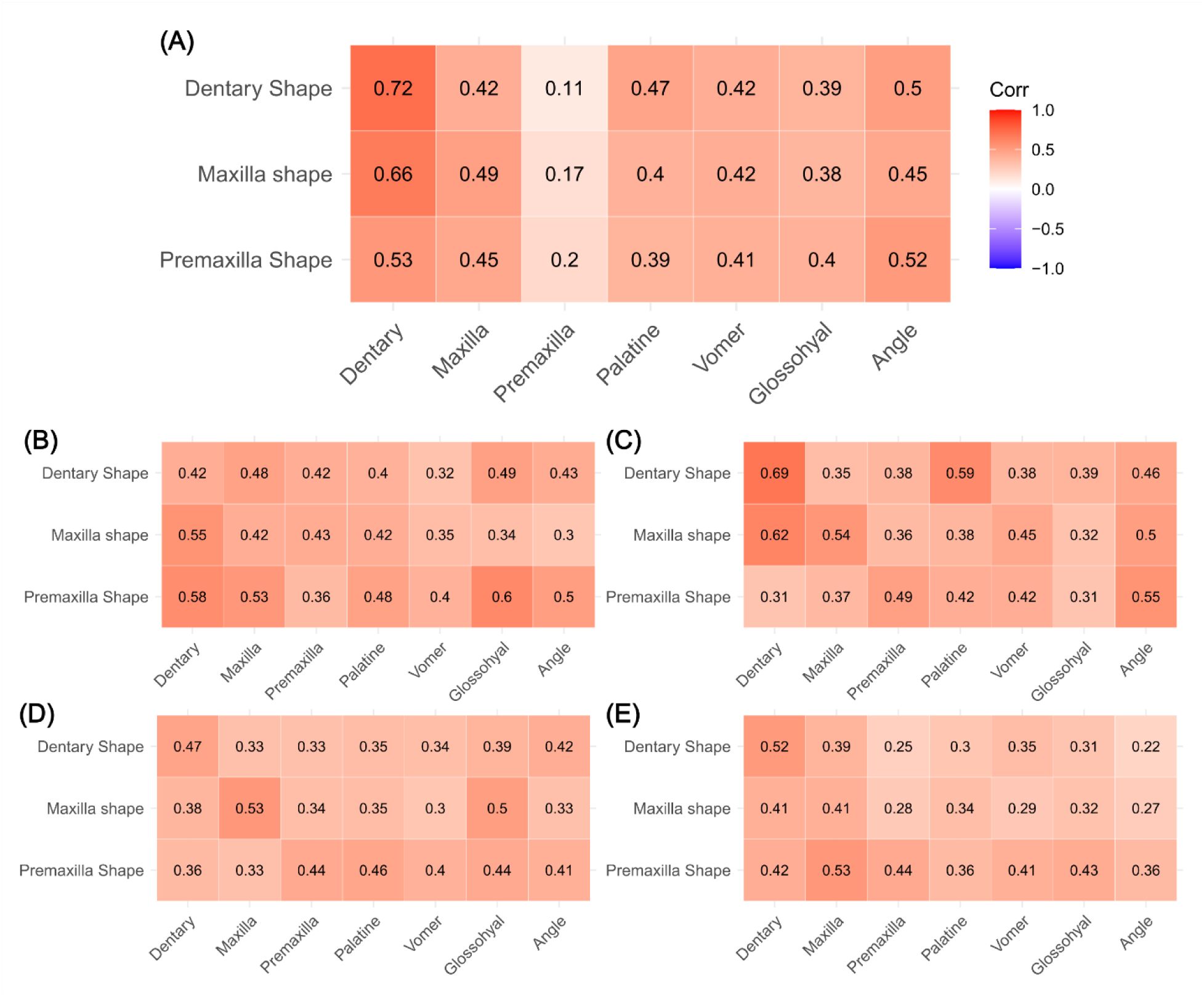
Heatmaps showing covariation between bone shape and tooth traits (numbers and angle) by morph. Traits were compared using partial least squares (PLS), within and among (All) morph. Plot shows the estimated r-PLS. Panels show five correlations plot, A) all morph, B) PI-charr, C) PL-charr, D) LB-charr, E) SB-charr. See Online Resource 42 for further results on the covariation.

## Discussion

In general fishes tend to have uniform-shaped teeth and high intraspecific variation in tooth numbers (Gidmark et al., 2019). We explored variation in tooth numbers and inclination angle among sympatric Arctic charr morphs in lake Þingvallavatn, known for extensive variation in external head and jaw bone morphology which appears to relate to their ecological specializations and life history characteristics (Jonsson et al., 1988; Jónsdóttir et al., 2024; Malmquist, 1992; Sandlund et al., 1987; Skúlason, Noakes, et al., 1989; Snorrason et al., 1989). Our data reveal variance for tooth numbers in six jaw bones (dentary, maxilla, premaxilla, palatine, vomer and glossohyal), and fluctuating asymmetry in the four paired bones. Tooth numbers on all bones increased with the size of the fish, except the premaxilla and glossohyal. We also found significant differences in tooth number between morphs in four bones (dentary, palatine, vomer and glossohyal), with pelagic morphs having more teeth than benthic morphs in all cases. There was a significant difference in maxilla tooth angle, with benthic morphs having more inward angled teeth. We found significant covariations between these tooth traits and bone shape.

### The curious lack of variance and allometry in premaxilla teeth counts

As most (but not all) fish show asymptotic growth related to maturity (Beverton & Holt, 1959), one expects their features would change with size, and some at different rate. We assumed that tooth numbers would relate positively with body size for all bones, i.e. more teeth as the fish grows. The confirmed this for all bones, except the premaxilla and glossohyal. These relationships did not differ by morph, not even for the dentary, where the shape-size allometry differs markedly by morph (Jónsdóttir et al., 2024). No relationship was found between size and maxilla tooth angles. We hypothesize three simple models for the relationship between tooth number and body size: 1) Tooth numbers are fixed and don’t change at all with size (flat line). 2) Tooth numbers increase with size to a specific number or body size at which point the number becomes fixed. 3) Tooth numbers increase with size. Either models 1 or 2 may apply to the glossohyal and the premaxilla and likely model 3 applies for the other bones. Since we did not sample juvenile fish, we cannot distinguish between model 1 and 2.

The premaxilla tooth numbers are not only stable over size, but also between morphs and within individuals. Fluctuating asymmetry is believed to stem from developmental noise generated by chance events (Koeberle et al., 2020; Merilä & Björklund, 1995; Van Valen, 1962). We found fluctuating asymmetry in tooth numbers for all four paired bones studied here. This suggests substantial developmental noise either during tooth formation or tooth replacement, in Arctic charr. The premaxilla had the most individuals with the same number of teeth on the left and right bone (no asymmetry), though only around 21%. It is interesting that this bone with the “least” fluctuating asymmetry also showed the least tooth number variation between individuals, no relationship between body size and tooth number, and no variation between morphs.

The relative stability of the premaxilla tooth numbers with size and the lack of differences between morphs confirms earlier results (Bryce et al., 2016; Ingimarsson, 2002). Could this lack of variation in tooth numbers relate to the relatively few teeth (4 to 10 in this study) on the premaxilla? With few teeth, an offset of one or two teeth could inflate the variance and reduce statistical power to estimate an allometric slope. However, the vomer has a similar number of teeth (1 to 9) yet showed a significant relationship with size and varied by morph. Therefore, we entertain the idea that the invariance of premaxilla tooth numbers in Arctic charr adults is biological. This could relate to a finite number of dental primordia being set early in premaxilla development, while in bones that gain teeth with increasing size, a stem cell niche may provide new precursor cells throughout the growth phase. Studies have shown that in other salmonid (rainbow trout (*Oncorhynchus mykiss*) both the timing and position of replacement teeth is regulated, with the genetic components which regulate the timing and position being local to each tooth family (i.e. a tooth and all of its successors, Fraser et al., 2006). The idea that dental diversity can arise by shifts in the replacement programs (Tucker & Fraser, 2014), was supported by studies on three-spined sticklebacks (Cleves et al., 2014; Ellis et al., 2015). Here we hypothesize that the tooth formation in the premaxilla teeth may be more regulated than the tooth formation of the other bones. Developmental constraints on premaxilla teeth numbers could explain our observations. Variation in the constraints on tooth number by bones calls for comparative analyses of the developmental programs involved.

Could this invariance in the premaxilla be a conserved trait among fishes and shared by other vertebrates? Perhaps not, as premaxilla tooth number increase with size in the three-spined stickleback, a relationship that differs between populations (Caldecutt et al., 2001). However, invariance of premaxilla tooth numbers was found in some tetrapods, for examples the reptiles *Ctenotus essigtoni* (Greer, 1991), American alligator (*Alligator mississippiensis*, Brown et al., 2015) and Komodo dragon (*Varanus komodoensis*, Brown et al., 2015). Analyses of fossilized *Tyrannosaurid* dinosaurs indicate that premaxilla tooth numbers were comparably stable within that group; not varying between taxa, individuals within a genus and with no asymmetry, though tooth numbers on other bones (maxilla and dentary) did (P. Currie, 2003; P. J. Currie, 2003). We add the caveat that even though premaxilla tooth numbers do not seem to vary between Arctic charr morphs (Bryce et al., 2016; Ingimarsson, 2002) other aspects of the premaxilla teeth (e.g. length, thickness, etc) could vary (Bryce et al., 2016). We call for further study of the allometries of teeth in more populations and species of salmonids.

### Differences in tooth traits between morphs within species

Based on prior studies (Bryce et al., 2016; Ingimarsson, 2002; Jónsdóttir et al., 2024) we predicted morph differences in tooth number in the dentary and palatine. The data confirmed this and indicated tooth numbers in the maxilla and premaxilla do not differ among morphs. The data also revealed morph differences in the two bones forming the “tongue-bite” apparatus (vomer and glossohyal). Differences between Arctic charr morphs have previously been documented for the dentary, palatine (Bryce et al., 2016; Ingimarsson, 2002) and the glossohyal (Pichugin, 2009), but not the vomer. Confirming Ingimarsson (2002) findings, we report morph differences in the inward inclination of maxillary teeth, the angle being larger in the benthic morphs. The pelagic morphs tended to have more teeth than the benthic morphs (in the dentary, palatine, vomer and glossohyal). This agrees with studies of adult Scottish Arctic charr and juvenile Siberian Arctic charr (Bryce et al., 2016; Pichugin, 2009). The differences in teeth numbers across bones may arise with variation in dental palette lengths, tooth replacement programs or other factors (Cleves et al., 2014; Ellis et al., 2015; Tucker & Fraser, 2014). Assuming that the ancestor of the morphs was mostly likely a piscivore (Bengtsson et al., 2023), one could postulate that the reduced teeth numbers in the benthic morphs is a derived trait.

As the estimated differences of tooth number are small (1 - 3 teeth) the question arises, how biologically important are they? With discrete characters like teeth and most bones having 10 or more teeth, addition or loss of one tooth may not affect fitness. Fishes, including salmonids, use teeth for capturing and holding prey and some species use them to process prey into consumable parts (Camp et al., 2009; Gidmark et al., 2019; Konow et al., 2008). The functional significance of these differences in tooth number remains unclear (Bryce et al., 2016), and they may not be adaptative. We previously found strong differences in the shape of the dentary bone between the Þingvallavatn morphs (Jónsdóttir et al., 2024) and hypothesized that shorter dentary (most likely the derived state) may aid benthic morphs in catching/holding their prey, from the stony littoral lake bottom. Here, the dentary tooth numbers correlated with the dentary shape. Thus, it’s possible that fewer dentary teeth reflect a correlated response to the adaptive bone shape change (i.e. reduced dental palate resulting in less teeth). This may also apply for the other bones (palatine, vomer and glossohyal), however we have not studied their shapes at this point (Jónsdóttir et al., 2024). It is also possible that the tooth number divergence is due to genetic drift, loss of purifying selection or genetic correlation as morphs are known to differ genetically (Guðbrandsson et al., 2019; Xiao et al., Unpublished) and in many other phenotypic traits (Jónsdóttir et al., 2024; Skúlason, Noakes, et al., 1989; Snorrason et al., 1989).

However, if these variations are adaptive then more teeth on the dentary and the palatine may make it easier to either grab or hold elusive prey. More teeth on both the vomer and glossohyal in pelagic morphs could reflect adaptions for prey-processing, as both bones are part of the tongue-bite apparatus in salmonids (Camp et al., 2009; Konow et al., 2008). Stomachs of LB- and SB-charr contain primarily whole snails (Malmquist, 1992), which suggests they rarely process their prey before swallowing. However, having more teeth on the same bones used for processing prey by other piscivorous salmonids (Konow et al., 2008; Konow & Sanford, 2008) may allow PI-charr to consume relatively large-bodied prey by either increasing handling efficiency or by allowing prey to be broken down into smaller consumable parts (Konow et al., 2008). If increased tooth count aids processing efficiency, it may also explain why larger fish have more teeth as they generally eat larger prey. Processing efficiency has not been directly studied in this system. The increased tooth numbers on the same bones in the PL-charr, which primarily eats copepods (Malmquist, 1992), however, does not have a clear ecological benefit. Little is known of the raking behaviour in the Þingvallavatn charr morphs, and within the *Salvelinus* genus, modulation of muscle activity or raking kinematics of the tongue-bite apparatus based on prey type is little studied (Konow et al., 2008). Therefore, we cannot confirm that the tooth differences in the vomer and glossohyal reflect adaptive differences in prey-processing.

While the tooth number variations between morphs may not be adaptive, there is a possibility that the variation in maxillary tooth angle could be. The strongest difference in tooth angles was between the smaller morphs (SB- and PL-charr), not the larger two (LB- and PI-charr) which differed mainly in tooth numbers in other bones. SB-charr had an average angle of 50.9°, while the PL-charr 32.4° (in between were LB-charr: 48.3° and PI-charr: 40.8°). Variation in tooth angle is understudied in salmonids and teleosts. Anatomical species descriptions indicate backwards curvature of glossohyal teeth in salmonids (Day, 1887) and inward curvature of premaxilla teeth in predaceous catfishes (Gosline, 1973), and one hypothesis is that this prevents prey from escaping the mouth (Gosline, 1973). It is unlikely that the inward pointing maxilla teeth in SB-charr serve a similar role, as snails (their main prey) are not particularly evasive prey. However, in the lesser electric ray (*Narcine brasiliensis*) angle teeth are believed to hold the prey in place while sediment is cleaned and expelled (Dean et al., 2008). It is possible that angled teeth of the benthic charr morphs serve a similar role, preventing the prey from being swallowed before sediment is cleaned. The angled teeth may also assist in holding the prey or orienting it for swallowing. Ingimarsson (2002) proposed the inward angled teeth may allow benthic morphs to hook their teeth underneath snails from the lava bottom and dislodge them. This “soda-bottle opener” hypothesis, and others listed above, could be tested with observations of feeding fish. Studies of more snail eating charr morphs can evaluate the generalizability of these findings. We also note the analyses suggested differences between angles of the 1^st^ and 5^th^ tooth, with the 5^th^ tooth being more angled inwards in the larger morphs, but the 1^st^ tooth leaning more inwards in the smaller morphs. However, these differences in angle by tooth were minor. Examining variation in tooth angles in multiple teeth may be another interesting research avenue.

Other aspects of teeth not investigated here, like tooth length, width, pointedness and overall shape, could also be biologically meaningful (Karagic et al., 2020). These traits are known to differ among fish species (Gidmark et al., 2019; Streelman et al., 2003) and within Arctic charr (Bryce et al., 2016). For example, a long-standing proposition has been that piscivorous salmonids have larger teeth than non-piscivores (Day, 1887). Indeed, Bryce et al. (2016) reported piscivorous Arctic charr morph had longer teeth on the premaxilla, maxilla, dentary and glossohyal (Bryce et al., 2016). As sympatric polymorphism is widespread among Arctic charr (Doenz et al., 2019; Skoglund et al., 2015; Woods et al., 2013; Østbye et al., 2020), it is possible that many other Arctic charr populations show similar differences in tooth traits.

### Methodological aspects and future directions

Lastly, we discuss dataset differences, methodological limitations and suggest future directions. We analysed two datasets and despite qualitatively similar results, there were quantitative discrepancies (mainly fewer statistically significant terms in the “second dataset”). This could reflect differences in sample size, photography, layout of bones (3D structures – photographed as 2D), measures of traits and biological reality. For both datasets we applied the same reasoning for counting teeth, though the image quality was worse for the “second dataset” and two different researchers counted each set. Generally, there were minor quantitative differences in tooth counts between sets, most likely reflecting different sample sizes (power of statistical analysis), size distribution of samples (can affect tooth number, see also (Cleves et al., 2014; Ellis et al., 2015; Greer, 1991; Torres-Carvajal, 2007) and sampler effects (minor differences in tooth count due to non-identical humans). The notable exception was the glossohyal, ∼7 more teeth were found in the primary dataset. This most likely reflects the poor quality of the photos from 1997, causing tooth numbers to be underestimated. There were notable differences between datasets in the maxilla tooth angles, with more inwards leaning teeth in the primary dataset. This likely reflects differences in approaches to estimated tooth angles. For the primary dataset we measured angles with the medial/ventral method, as we found the fronto-lateral method (used for the “second dataset”) was not as repeatable (see above). Related to this, we measured the 1^st^ and 5^th^ teeth, but in the “second dataset” 2^nd^-5^th^ tooth was measured, depending on focus. The indication of angle differences by tooth could influence this as well. However, it is possible that tooth numbers and maxilla angles have changed in these morphs in the 20+ years between the two samplings.

We explored asymmetry and found it quite substantial for both tooth number and maxillary tooth angle in adult spawning individuals. Previous research has shown that asymmetry can vary between pre-spawning and post spawning salmon (Witten et al., 2005). Whether such patterns also hold in Arctic charr is unknown and was not tested here. Neither did we analyse asymmetry for the bone shapes. In the future, a joint analysis of asymmetry in bone shape, size and tooth traits, in more groups (with larger sample sizes) would be an exciting research avenue. Related to this are analyses of covariation of traits (not their asymmetry), bone shape and tooth numbers. We anticipated a relationship of tooth number and bone shape (Cleves et al., 2014; Ellis et al., 2015) and found support for this in the dentary and maxilla (not premaxilla) when analysed per morph. The analyses of the overall dataset indicated significant covariation of bone shape of the dentary, maxilla and premaxilla and tooth numbers on all the bones (except for premaxilla teeth). We think this stems from the landmark data retaining aspects of size (both body and bone size differ by morph, (Jónsdóttir et al., 2024; Sandlund et al., 1992), bone shape and/or tooth number differences by morph. Statistical power may also have influenced these results, the total samples size was 240 specimens. With larger sample sizes it would be interesting to estimate if overall bone shape correlates with tooth traits in other bones (palatine, vomer and glossohyal), and whether tooth numbers relate strongly with specific axes (e.g. elongation of dental palate) of bone shape in charr and other salmonids. Previous research on three-spined sticklebacks found a positive correlation between tooth plate areas and tooth number on pharyngeal jaw (Cleves et al., 2014; Ellis et al., 2015). Something similar may be occurring in the dentary, where pelagic morphs with more elongated dental palates, (Jónsdóttir et al., 2024) and a higher dentary tooth number. Differences in strength of correlation of tooth numbers and dental palate length in specific bones (or populations) could reflect developmental or evolved differences in the pathways responsible for tooth formation and growth.

## Conclusions

Our study shows that tooth traits (both their numbers and angles on maxilla) vary between recently diverged sympatric Arctic charr morphs. We found significant morph differences in four bones, three of which were previously shown to differ in other Arctic charr morphs (Bryce et al., 2016; Pichugin, 2009) and were able to confirm a prior finding on maxilla tooth angle differences (Ingimarsson, 2002). The functional or evolutionary relevance of these variations remain unknown and calls for further comparative work and behaviour surveys. Though we hypothesize that the tooth angle differences may be adaptive, with more angled teeth assisting the benthic morphs in either grabbing (Ingimarsson, 2002) or holding their prey (Dean et al., 2008). Although the allometry of tooth traits (both number and angle) did not differ between morphs it curiously varied markedly by bone. While some bones (e.g. dentary and palatine) showed strong allometry, others (e.g. premaxilla number and maxilla angle) did not. In our study it appears that the premaxilla dentition is more stereotypical than the other bones. Which may be due to either the development of tooth formation or replacement being more canalized in that bone. What does this say about the development of tooth formation on these diverse bones, and how and when during vertebrate evolution did development of particular structures become more stereotypical (and not affected by variation in size)? The great diversity in feeding and niche utilization in Arctic charr provides a way to investigate the developmental origins of adaptive and non-adaptive divergence of bone shape and dental variation.

## Supporting information

Combined supplemental files

## Acknowledgements

This study was financially supported by the Icelandic Research Fund, Ph.D. grant no. 2410464-051 to GOJ. We thank members of the Arctic charr group at the University of Iceland for their help during the project, especially Anthony Curtat for measuring tooth angle, Laura-Marie von Elm, Samuel Tersigni and Nahal Eskafi for assisting with processing and aging of fish. We thank heartfully Jóhann Jónsson and Rósa Jónsdóttir, farmers at Mjóanes, who both passed away recently, for their kind on-site support for this project. We thank Ian M. Dworkin for advice on the statistical analysis.

## Ethical Standards

This study builds on a previous study, and all the biological samples were ready in house. The previous study involved sacrificing of wild fishes, and for such work scientific fish fieldwork from the Directorate of Fisheries in Iceland (www.fiskistofa.is) was needed. Sigurður S. Snorrason and Arnar Pálsson, authors on both of these studies and in charge and present for all sampling efforts and, had for the period of the study, such scientific fish fieldwork permits (#0042/2014-2.13 and #0460/2021-2.0). The samples from 1999 were gathered before this law came into effect and are thus exempt from its stipulations.

## Supporting Information

***Online Resource 1*** *Morphology of two maxilla bones from three different views. Example of fronto-lateral view (A and B) ventral view (C and D) and medial view (E and F) images used to measure/calculate tooth angle. Shown are bones of individuals (A) PC3840 (male LB-charr, length = 37.1 cm, weight = 672 g), (B, D and F) PC3907 (female SB-charr, L = 15 cm, W = 24.5 g) and (C and E) PC3859 (male PL-charr, L = 14.8 cm, W = 36 g). Scale inset per picture is 1 cm*.

***Online Resource 2*** *Linear regression was used to estimate repeatability of tooth counts for the A) dentary, B) maxilla, C) premaxilla, D) palatine, E) vomer and F) glossohyal. Scatterplot shows the relationship between the first (March) and second (May) counts. Equations for each line are provided in Online Resource 3*.

***Online Resource 3*** *The results of liner regression models (using function lm) between the two teeth counts (March and May). Visualization of the datapoints and regression line are provided in Online Resource 2*.

***Online Resource 4*** *Repeatability of tooth counts analysed with Pearson correlation (function: cor.test), for each morph separately. Table shows the estimate (r) and results from the test of correlation between the two teeth counts (March vs May). For paired bones the correlation was tested for both the left side and right side separately, and the average number of teeth over both sides. Only one estimate was done for the non-paired bones (vomer and glossohyal)*.

***Online Resource 5*** *Fluctuating asymmetry in tooth numbers on four paired bones in four sympatric Arctic charr morphs. Graphed are histograms of the within individual variation in tooth number by morph. The distributions of deviation in tooth numbers between the sides (calculated by subtracted right from left), for the dentary (A), maxilla (B), premaxilla (C) and palatine (D). X-axis shows the deviation between left and right and the Y-axis the number of individuals per bin. Note the scale is continuous, as represented are the average from two measures per individual*.

***Online Resource 6*** *Distribution of the repeated measures for each of the paired bones, for each morph. The scatterplots show the linear relationship between tooth numbers from the right and left bones (A: dentary, B: maxilla, C: premaxilla and D: palatine). Equations for each line can be seen in Online Resource 7*.

***Online Resource 7*** *Results of linear regression model between tooth numbers of the right and left bones for all paired bones (dentary, maxilla, premaxilla and palatine). For visualization of relevant data, see Online Resource 6*.

***Online Resource 8*** *Results from student t-test, testing whether the mean asymmetry in tooth numbers differs from zero. Bonferroni corrected threshold (α = 0.01) to correct for tests on four bones*.

***Online Resource 9*** *Test of the influence of size (ln fork length), morph and the interaction on fluctuating asymmetry in tooth numbers in the four symmetric bones. Results from GLM with ln FL for size, the morphs (four types) and the interaction term*.

***Online Resource 10*** *The scatterplots show deviation in tooth numbers between left and right (Y-axis) against size (ln FL in cm, X-axis) for each of the four paired bones, dentary (A and B), maxilla (C and D), premaxilla (E and F) and palatine (G and H). The lines are a fitted linear model, with confidence intervals, nonsignificant by ANOVA on GLM models. Graphs on right (A, C, E and G) show all morph together and graphs on left (B, D, F and H) are coloured by morph*.

***Online Resource 11*** *Correlations (Pearson) between mean asymmetry in tooth numbers for each bone to each other and fork length*.

***Online Resource 12*** *Average number of teeth for six bones and respective standard deviation (SD), calculated for all Arctic charr morphs. The numbers represent raw counts*.

***Online Resource 13*** *Distribution of tooth number in six bones by morph. Also shown are sex specific distributions as boxplots. The Y-axis indicates the number of teeth (average per individual for the paired bones A-D, average over repeated measures for the non-paired E-F). The bones are: dentary (A), maxilla (B), premaxilla (C), palatine (D), vomer (E) and glossohyal bone (F). Morphs are coloured: SB: dark blue, LB: light blue, PI: orange, PL: red (see also inset). Superimposed are boxplots for the sexes (female: purple, males: green, not sexed: grey, only in SB- charr)*.

***Online Resource 14*** *The size corrected distribution of tooth number in six bones by morph. Also shown are sex specific distributions as boxplots. The Y-axis indicates the number of teeth corrected for with individual length (FL) (teeth/length). The bones are: dentary (A), maxilla (B), premaxilla (C), palatine (D), vomer (E) and glossohyal bone (F). Morph are coloured: SB: dark blue, LB: light blue, PI: orange, PL: red (see also inset). Superimposed are boxplots for the sexes (females: purple, males: green, not sexed: grey, only in SB-charr)*.

***Online Resource 15*** *Result from emtrends, showing the different trends between tooth number and different length measurements (raw fork length, ln fork length, ln C-size)*.

***Online Resource 16*** *The influence of size, morph, sex and side (left and right) on tooth number in six jaw bones. Analysed with generalized liner mixed model (glmmTMB) using ln C-size for size*.

***Online Resource 17*** *Pairwise comparisons of teeth numbers between morphs, results from the generalized liner mixed models (glmmTMB), when using ln FL for size*.

***Online Resource 18*** *Pairwise comparisons of teeth numbers between morphs within sex, results from the generalized liner mixed models (glmmTMB), when using ln FL for size*.

***Online Resource 19*** *Pairwise comparisons of teeth numbers between sexes within morph, results from the generalized liner mixed models (glmmTMB), when using ln FL for size*.

***Online Resource 20*** *Pairwise comparisons of teeth numbers between morphs, results from the generalized liner mixed models (glmmTMB), when using ln C-size for size*.

***Online Resource 21*** *Pairwise comparisons of teeth numbers between morphs within sex, results from the generalized liner mixed models (glmmTMB), when using ln C-size for size*.

***Online Resource 22*** *Pairwise comparisons of teeth numbers between sexes within morph, results from the generalized liner mixed models (glmmTMB), when using ln C-size for size*.

***Online Resource 23*** *Linear regression (function lm) examining the repeatability of the two methods (fronto/lateral method and medial/ventral method) used when measuring the angle of maxillary teeth. This was analysed on 62 specimens from 3 morphs. The Medial/ventral method appeared to be more repeatable*.

***Online Resource 24*** *Test of repeatability for estimation of maxillary teeth angle with linear regression. Scatterplots show the liner relationship between the two trials for both methods (fronto/lateral method and medial/ventral method) used to measure the angle. This was analysed on 62 specimens from 3 morphs. The Medial/ventral method appeared to be more repeatable*.

***Online Resource 25*** *Average angle of maxillary teeth (*± *standard deviation) for each morph, also by sex*.

***Online Resource 26*** *Pairwise comparisons of the inward inclination of maxillary teeth angles between morphs, results from the generalized liner mixed models (glmmTMB), when using ln FL for size*.

***Online Resource 27*** *Pairwise comparisons of the inward inclination of maxillary teeth angles between morphs within sex, results from the generalized liner mixed models (glmmTMB), when using ln FL for size*.

***Online Resource 28*** *Pairwise comparisons of inward inclination maxillary teeth angles between sexes within morph, results from the generalized liner mixed models (glmmTMB), when using ln FL for size*.

***Online Resource 29*** *The influence of size, morph, sex, side (left or right) and tooth (1^st^ or 5^th^ tooth) on the inward inclination of maxillary teeth. Analysed with generalized liner mixed model (glmmTMB) using ln C-size for size*.

***Online Resource 30*** *Pairwise comparisons of the inward inclination of maxillary teeth angles between morphs, results from the generalized liner mixed models (glmmTMB), when using ln C-size for size*.

***Online Resource 31*** *Pairwise comparisons of the inward inclination of maxillary teeth angles between morphs within sex, results from the generalized liner mixed models (glmmTMB), when using ln FL for size*.

***Online Resource 32*** *Pairwise comparisons of the inward inclination of maxillary teeth angles between sexes within morph, results from the generalized liner mixed models (glmmTMB), when using ln FL for size*.

***Online Resource 33*** *Effects of morph and sex on the inward inclination of maxillary teeth in the four morphs. The scatterplot shows average tooth angle (averaged over 1^st^ and 5^th^ tooth on both left and right bone for each individual) (Y-axis) against size (fork length of fish, X-axis). The ellipses and shaded areas are 95% CI for the distributions and the large dots represent the means for the sexes within morphs*.

***Online Resource 34*** *Average number of teeth for six bones and respective standard deviation (SD), calculated for all Arctic charr from the 1997 “replicate” dataset. The numbers represent raw counts*.

***Online Resource 35*** *Distribution of tooth number in six bones by morph, based on the 1997 “replicate” dataset. Also shown are sex specific distributions as boxplots. The Y-axis indicates the number of teeth. The bones are: dentary (A), maxilla (B), premaxilla (C), palatine (D), vomer (E) and glossohyal bone (F). Morphs are coloured: SB: dark blue, LB: light blue, PI: orange, PL: red (see also inset). Superimposed are boxplots for the sexes (female: purple, males: green)*.

***Online Resource 36*** *The influence of size, morph and sex on tooth number in six jaw bones, from the 1997 “replicate” dataset. Analysed with generalized liner mixed models (glmmTMB) using ln FL for size. P-values below the Bonferroni corrected threshold (α = 0.008) are shown in bold*.

***Online Resource 37*** *Influence of size and morph on number of teeth from the 1997 “replicate” dataset. The scatterplots show individual tooth numbers (Y-axis) against size (ln FL in cm, X-axis) for the four morphs and six bones; A) dentary, B) maxilla, C) premaxilla, D) palatine, E) vomer and F) glossohyal. Each plot shows p-values from glmmTMB for the effect of morph (M), body size (FL) and morph by body size (MxFL) on tooth number*.

***Online Resource 38*** *Average angle of maxilla teeth (*± *standard deviation) for each morph, also by sex from the 1997 “replicate” dataset*.

***Online Resource 39*** *The influence of size, morph and sex on the inward inclination of maxillary teeth, from the 1997 “replicate” dataset. Analysed with generalized liner mixed models (glmmTMB) using ln FL for size*.

***Online Resource 40*** *Maxilla tooth angle from the 1997 “replicate” dataset varies by morphs (A) and sex by morph (B). The scatterplots show average teeth angle for 1^st^ and 5^th^ tooth on both left and right bone (Y-axis) against size (Fork length of fish, X-axis). Maxilla teeth angle varies by morphs (A) and sex by morph (B). The ellipses and shaded areas are 95% CI for the distributions and the large dots represents the mean for each group. Two datasets were analysed, 240 fish from Jonsdottir 2024, combined data for tooth 1 and 5, with medial/ventral method (see figure 5 and Online Resource 33) and 138 fish from Ingimarsson 2001, measured on tooth 2-5 with fronto-lateral method*.

***Online Resource 41*** *Test of integration between bone shape and tooth number. Traits were compared using partial least squares (PLS), within and among bones*.

***Online Resource 42*** *Test of covariation between bone shape and tooth traits (numbers and angle) by morph. Traits were compared using partial least squares (PLS), within and among (All) morphs. Corresponding pairs of tooth counts and shape are underlined*.

