## Supplementary material for "Variation of tooth traits in ecologically specialized and sympatric morphs": Combined supplemental files

For manuscript

Submitted to Evolutionary Biology

Authors: Guðbjörg Ósk Jónsdóttir[1*], Finnur Ingimarsson[1,2], Sigurður Sveinn Snorrason[1], Sarah Elizabeth Steele[3] and Arnar Pálsson[1]

ORCID:

GOJ: https://orcid.org/0009-0008-0502-5553

FI: https://orcid.org/0000-0002-0815-7622

SES: https://orcid.org/0000-0001-8404-5537

AP: https://orcid.org/0000-0002-6525-8112

Affiliations

1. Institute of Life- and Environmental Science, University of Iceland
2. Thingvellir national park, Iceland
3. Canadian Museum of Nature, Ottawa, Canada

Corresponding author:

* Guðbjörg Ósk Jónsdóttir,

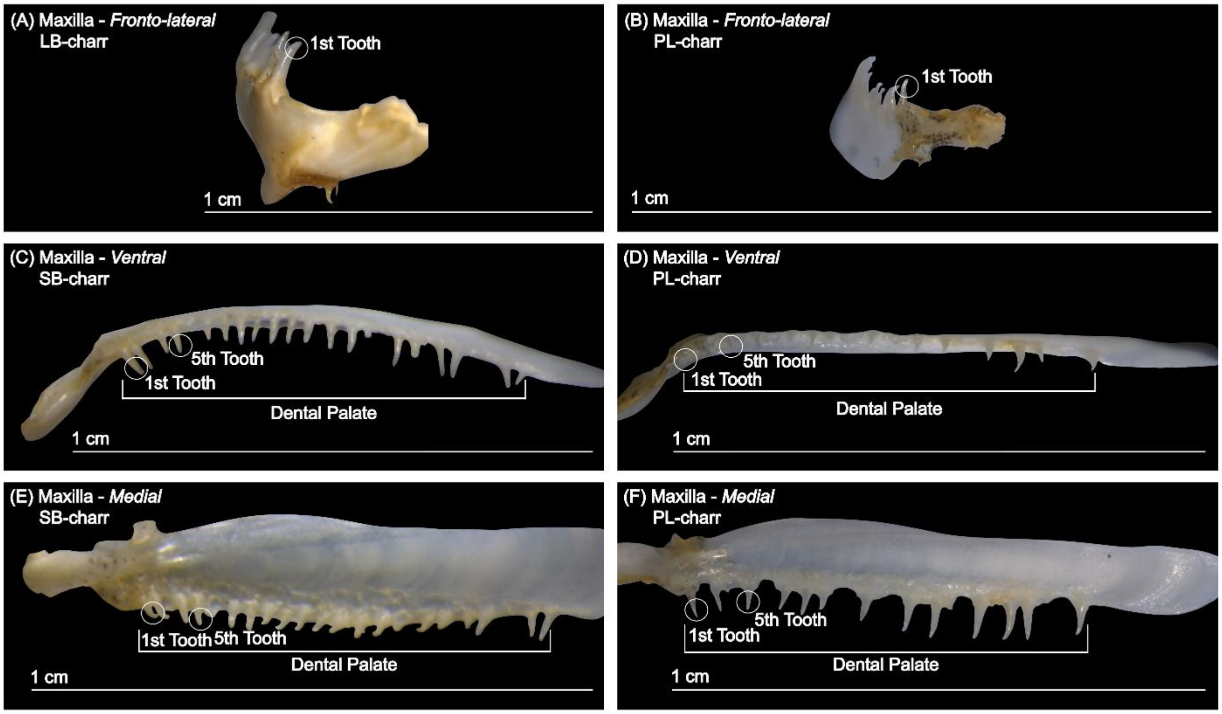


***Online Resource 1*** *Morphology of two maxilla bones from three different views. Example of fronto-lateral view (A and B) ventral view (C and D) and medial view (E and F) images used to measure/calculate tooth angle. Shown are bones of individuals (A) PC3840 (male LB-charr, length = 37.1 cm, weight = 672 g), (B, D and F) PC3907 (female SB-charr, L = 15 cm, W = 24.5 g) and (C and E) PC3859 (male PL-charr, L = 14.8 cm, W = 36 g). Scale inset per picture is 1 cm.*

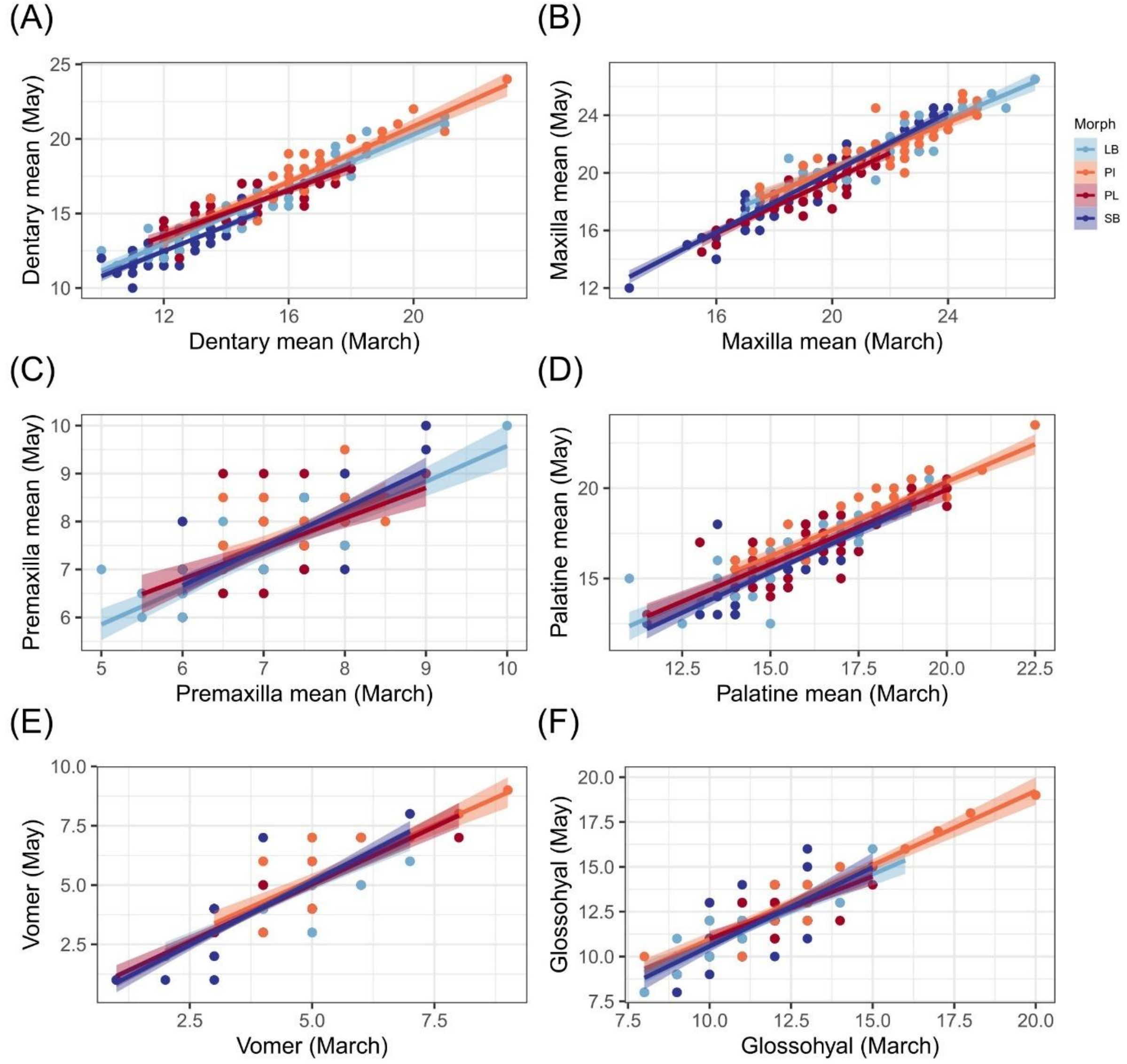


***Online Resource 2*** *Linear regression was used to estimate repeatability of tooth counts for the A) dentary, B) maxilla, C) premaxilla, D) palatine, E) vomer and F) glossohyal. Scatterplot shows the relationship between the first (March) and second (May) counts. Equations for each line are provided in Online Resource 3.*

***Online Resource 3*** *The results of liner regression models (using function lm) between the two teeth counts (March and May). Visualization of the datapoints and regression line are provided in Online Resource 2.*

| Bones | Side | Morph | $r^2$ | Slope | Intercept | p-value |
|---|---|---|---|---|---|---|
| Dentary | Mean | All | 0.888 | 0.99 | 0.78 | < 0.001 |
| | | PL | 0.727 | 0.77 | 4.29 | < 0.001 |
| | | PI | 0.851 | 0.93 | 2.25 | < 0.001 |
| | | LB | 0.897 | 0.91 | 2.02 | < 0.001 |

| | | | | | | |
|---|---|---|---|---|---|---|
| | | SB | 0.720 | 0.85 | 2.32 | < 0.001 |
| | Right | All | 0.846 | 0.95 | 1.30 | < 0.001 |
| | | PL | 0.675 | 0.69 | 5.36 | < 0.001 |
| | | PI | 0.821 | 0.83 | 3.57 | < 0.001 |
| | | LB | 0.828 | 0.96 | 1.16 | < 0.001 |
| | | SB | 0.675 | 0.82 | 2.47 | < 0.001 |
| | Left | All | 0.818 | 0.96 | 1.44 | < 0.001 |
| | | PL | 0.642 | 0.77 | 4.24 | < 0.001 |
| | | PI | 0.758 | 0.94 | 2.46 | < 0.001 |
| | | LB | 0.823 | 0.81 | 3.75 | < 0.001 |
| | | SB | 0.618 | 0.77 | 3.41 | < 0.001 |
| Maxilla | Mean | All | 0.886 | 0.97 | 0.57 | < 0.001 |
| | | PL | 0.805 | 0.93 | 0.82 | < 0.001 |
| | | PI | 0.757 | 0.82 | 3.83 | < 0.001 |
| | | LB | 0.848 | 0.86 | 3.20 | < 0.001 |
| | | SB | 0.902 | 1.04 | -0.71 | < 0.001 |
| | Right | All | 0.837 | 0.89 | 2.03 | < 0.001 |
| | | PL | 0.725 | 0.88 | 1.89 | < 0.001 |
| | | PI | 0.744 | 0.77 | 4.92 | < 0.001 |
| | | LB | 0.740 | 0.74 | 5.60 | < 0.001 |
| | | SB | 0.865 | 0.95 | 0.90 | < 0.001 |
| | Left | All | 0.833 | 0.96 | 0.69 | < 0.001 |
| | | PL | 0.764 | 0.89 | 1.76 | < 0.001 |
| | | PI | 0.588 | 0.82 | 3.94 | < 0.001 |
| | | LB | 0.850 | 0.91 | 2.34 | < 0.001 |
| | | SB | 0.799 | 0.98 | 0.35 | < 0.001 |
| Premaxilla | Mean | All | 0.554 | 0.74 | 2.25 | < 0.001 |
| | | PL | 0.374 | 0.63 | 2.99 | < 0.001 |
| | | PI | 0.452 | 0.68 | 2.77 | < 0.001 |
| | | LB | 0.689 | 0.75 | 2.13 | < 0.001 |
| | | SB | 0.600 | 0.80 | 1.85 | < 0.001 |
| | Right | All | 0.480 | 0.69 | 2.67 | < 0.001 |
| | | PL | 0.431 | 0.71 | 2.44 | < 0.001 |
| | | PI | 0.426 | 0.66 | 2.93 | < 0.001 |
| | | LB | 0.607 | 0.67 | 2.69 | < 0.001 |
| | | SB | 0.411 | 0.69 | 2.67 | < 0.001 |
| | Left | All | 0.498 | 0.68 | 2.68 | < 0.001 |
| | | PL | 0.720 | 0.58 | 3.33 | < 0.001 |
| | | PI | 0.707 | 0.57 | 3.59 | < 0.001 |
| | | LB | 0.649 | 0.71 | 2.38 | < 0.001 |

| | | | | | | |
|---|---|---|---|---|---|---|
| | | SB | 0.552 | 0.72 | 2.43 | < 0.001 |
| Palatine | Mean | All | 0.785 | 0.91 | 2.05 | < 0.001 |
| | | PL | 0.690 | 0.83 | 3.41 | < 0.001 |
| | | PI | 0.830 | 0.83 | 3.83 | < 0.001 |
| | | LB | 0.737 | 0.84 | 3.10 | < 0.001 |
| | | SB | 0.733 | 0.90 | 1.86 | < 0.001 |
| | Right | All | 0.771 | 0.89 | 2.30 | < 0.001 |
| | | PL | 0.716 | 0.84 | 3.17 | < 0.001 |
| | | PI | 0.806 | 0.76 | 5.13 | < 0.001 |
| | | LB | 0.676 | 0.94 | 1.49 | < 0.001 |
| | | SB | 0.740 | 0.87 | 2.34 | < 0.001 |
| | Left | All | 0.774 | 0.90 | 2.10 | < 0.001 |
| | | PL | 0.692 | 0.81 | 3.59 | < 0.001 |
| | | PI | 0.772 | 0.89 | 2.72 | < 0.001 |
| | | LB | 0.734 | 0.84 | 3.12 | < 0.001 |
| | | SB | 0.772 | 0.89 | 1.91 | < 0.001 |
| Vomer | | All | 0.817 | 0.99 | 0.11 | < 0.001 |
| | | PL | 0.803 | 0.97 | 0.20 | < 0.001 |
| | | PI | 0.696 | 0.92 | 0.62 | < 0.001 |
| | | LB | 0.821 | 0.91 | 0.40 | < 0.001 |
| | | SB | 0.777 | 1.06 | -0.19 | < 0.001 |
| Glossohyal | | All | 0.706 | 0.84 | 2.34 | < 0.001 |
| | | PL | 0.525 | 0.70 | 3.99 | < 0.001 |
| | | PI | 0.849 | 0.83 | 2.70 | < 0.001 |
| | | LB | 0.682 | 0.78 | 2.90 | < 0.001 |
| | | SB | 0.505 | 0.88 | 1.76 | < 0.001 |

r2 = coefficient of determination; p = p-values,

***Online Resource 4*** *Repeatability of tooth counts analysed with Pearson correlation (function: cor.test), for each morph separately. Table shows the estimate (r) and results from the test of correlation between the two teeth counts (March vs May). For paired bones the correlation was tested for both the left side and right side separately, and the average number of teeth over both sides. Only one estimate was done for the non-paired bones (vomer and glossohyal).*

| Bone | Side | Morphs | *r* | df | t-value | p-value |
|---|---|---|---|---|---|---|
| Dentary | Mean | All | 0.943 | 237 | 43.42 | < 0.001 |
| | | PL | 0.855 | 59 | 12.69 | < 0.001 |
| | | PI | 0.924 | 45 | 16.23 | < 0.001 |
| | | LB | 0.948 | 47 | 20.47 | < 0.001 |
| | | SB | 0.850 | 80 | 14.45 | < 0.001 |
| | Right | All | 0.920 | 236 | 36.05 | < 0.001 |

| | | | | | | |
|---|---|---|---|---|---|---|
| | | PL | 0.825 | 58 | 11.11 | < 0.001 |
| | | PI | 0.908 | 45 | 14.56 | < 0.001 |
| | | LB | 0.912 | 47 | 15.26 | < 0.001 |
| | | SB | 0.824 | 80 | 13.02 | < 0.001 |
| | Left | All | 0.905 | 237 | 32.68 | < 0.001 |
| | | PL | 0.805 | 59 | 10.43 | < 0.001 |
| | | PI | 0.873 | 45 | 12.03 | < 0.001 |
| | | LB | 0.909 | 47 | 14.96 | < 0.001 |
| | | SB | 0.789 | 80 | 11.49 | < 0.001 |
| Maxilla | Mean | All | 0.942 | 237 | 43.05 | < 0.001 |
| | | PL | 0.899 | 59 | 15.77 | < 0.001 |
| | | PI | 0.873 | 46 | 12.14 | < 0.001 |
| | | LB | 0.923 | 47 | 16.43 | < 0.001 |
| | | SB | 0.950 | 79 | 27.12 | < 0.001 |
| | Right | All | 0.915 | 231 | 34.55 | < 0.001 |
| | | PL | 0.854 | 57 | 12.39 | < 0.001 |
| | | PI | 0.866 | 46 | 11.74 | < 0.001 |
| | | LB | 0.863 | 46 | 11.60 | < 0.001 |
| | | SB | 0.931 | 76 | 22.27 | < 0.001 |
| | Left | All | 0.913 | 232 | 34.09 | < 0.001 |
| | | PL | 0.876 | 58 | 13.86 | < 0.001 |
| | | PI | 0.773 | 45 | 8.17 | < 0.001 |
| | | LB | 0.924 | 46 | 16.38 | < 0.001 |
| | | SB | 0.895 | 77 | 17.62 | < 0.001 |
| Premaxilla | Mean | All | 0.746 | 238 | 17.26 | < 0.001 |
| | | PL | 0.620 | 59 | 6.07 | < 0.001 |
| | | PI | 0.681 | 46 | 6.31 | < 0.001 |
| | | LB | 0.834 | 47 | 10.37 | < 0.001 |
| | | SB | 0.778 | 80 | 11.06 | < 0.001 |
| | Right | All | 0.694 | 223 | 14.41 | < 0.001 |
| | | PL | 0.669 | 55 | 6.67 | < 0.001 |
| | | PI | 0.662 | 46 | 5.99 | < 0.001 |
| | | LB | 0.785 | 45 | 8.49 | < 0.001 |
| | | SB | 0.647 | 71 | 7.15 | < 0.001 |
| | Left | All | 0.707 | 225 | 15.02 | < 0.001 |
| | | PL | 0.520 | 53 | 4.43 | < 0.001 |
| | | PI | 0.569 | 45 | 4.64 | < 0.001 |
| | | LB | 0.771 | 47 | 8.30 | < 0.001 |
| | | SB | 0.805 | 74 | 11.66 | < 0.001 |
| Palatine | Mean | All | 0.887 | 236 | 29.47 | < 0.001 |

| | | | | | | |
|---|---|---|---|---|---|---|
| | | PL | 0.834 | 59 | 11.60 | < 0.001 |
| | | PI | 0.913 | 46 | 15.17 | < 0.001 |
| | | LB | 0.861 | 47 | 11.63 | < 0.001 |
| | | SB | 0.858 | 78 | 14.75 | < 0.001 |
| | Right | All | 0.878 | 230 | 27.87 | < 0.001 |
| | | PL | 0.849 | 58 | 12.23 | < 0.001 |
| | | PI | 0.900 | 46 | 13.99 | < 0.001 |
| | | LB | 0.826 | 46 | 9.95 | < 0.001 |
| | | SB | 0.862 | 74 | 14.63 | < 0.001 |
| | Left | All | 0.880 | 227 | 27.97 | < 0.001 |
| | | PL | 0.835 | 54 | 11.15 | < 0.001 |
| | | PI | 0.881 | 46 | 12.65 | < 0.001 |
| | | LB | 0.860 | 46 | 11.43 | < 0.001 |
| | | SB | 0.880 | 75 | 16.05 | < 0.001 |
| Vomer | | All | 0.904 | 225 | 31.74 | < 0.001 |
| | | PL | 0.898 | 51 | 14.61 | < 0.001 |
| | | PI | 0.838 | 45 | 10.31 | < 0.001 |
| | | LB | 0.908 | 47 | 14.85 | < 0.001 |
| | | SB | 0.883 | 76 | 16.41 | < 0.001 |
| Glossohyal | | All | 0.841 | 234 | 23.79 | < 0.001 |
| | | PL | 0.730 | 59 | 8.20 | < 0.001 |
| | | PI | 0.923 | 46 | 16.31 | < 0.001 |
| | | LB | 0.830 | 46 | 10.08 | < 0.001 |
| | | SB | 0.715 | 77 | 8.98 | < 0.001 |

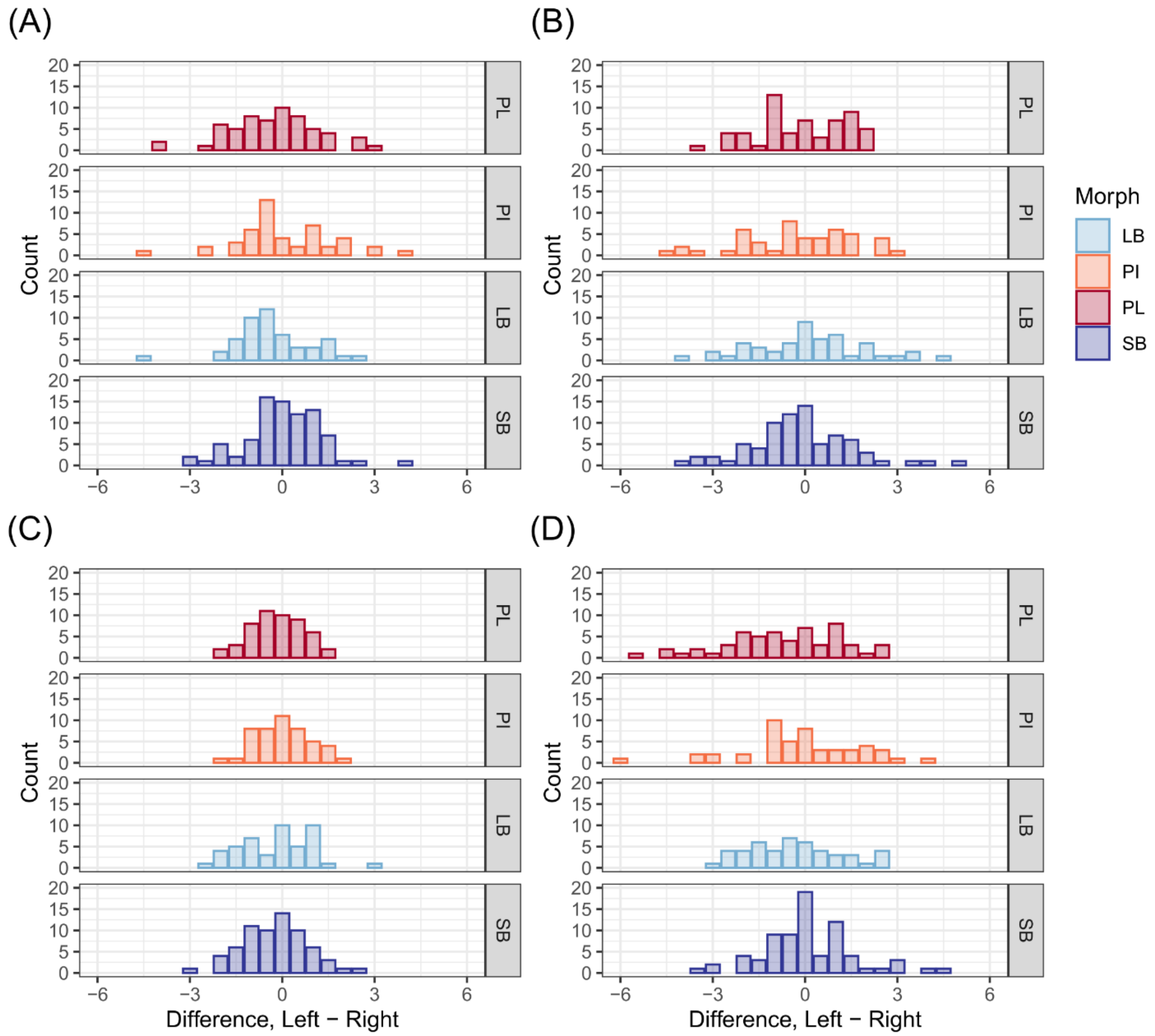


***Online Resource 5*** *Fluctuating asymmetry in tooth numbers on four paired bones in four sympatric Arctic charr morphs. Graphed are histograms of the within individual variation in tooth number by morph. The distributions of deviation in tooth numbers between the sides (calculated by subtracted right from left), for the dentary (A), maxilla (B), premaxilla (C) and palatine (D). X-axis shows the deviation between left and right and the Y-axis the number of individuals per bin. Note the scale is continuous, as represented are the average from two measures per individual.*

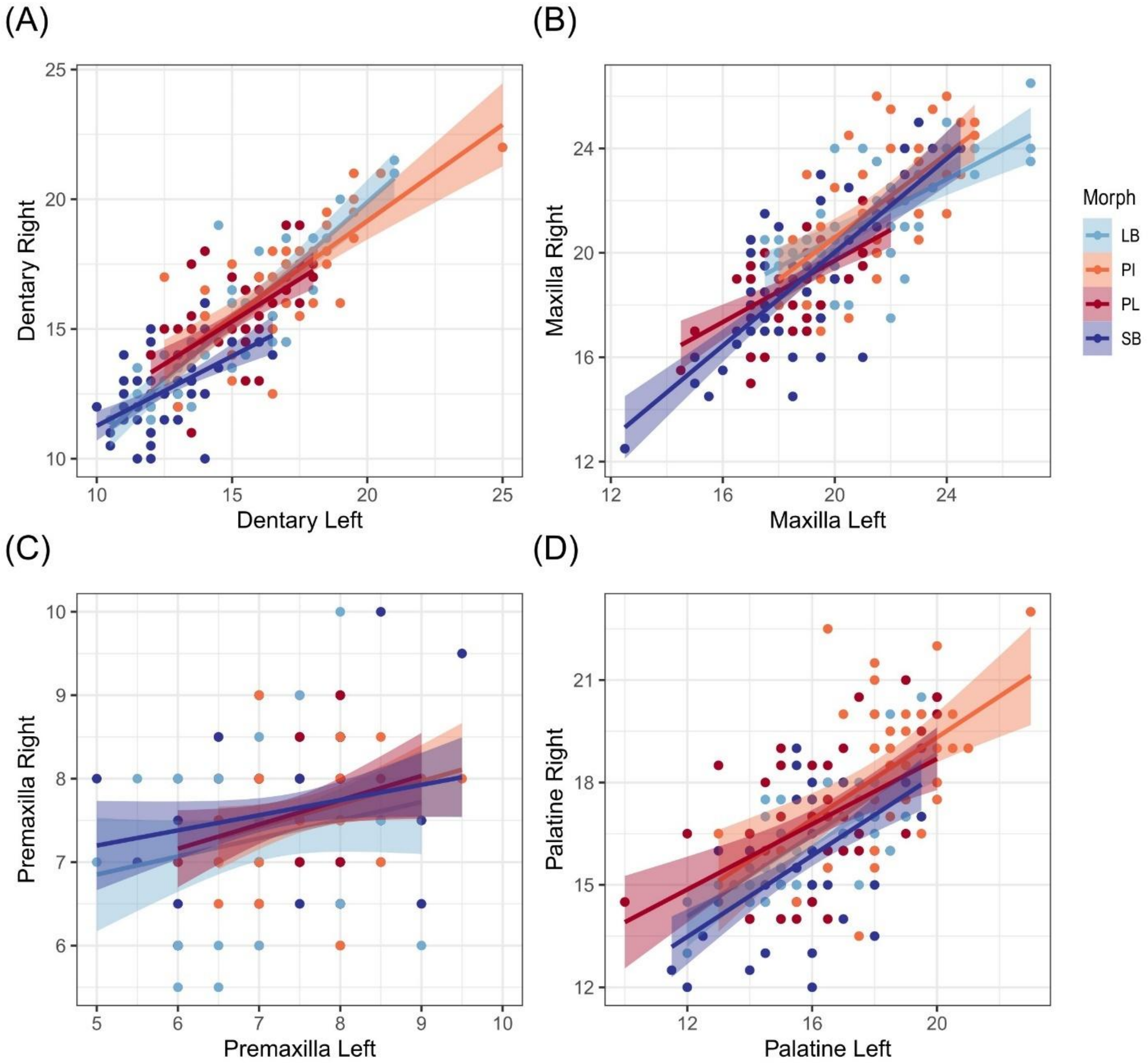


***Online Resource 6*** *Distribution of the repeated measures for each of the paired bones, for each morph. The scatterplots show the linear relationship between tooth numbers from the right and left bones (A: dentary, B: maxilla, C: premaxilla and D: palatine). Equations for each line can be seen in Online Resource 7.*

***Online Resource 7*** *Results of linear regression model between tooth numbers of the right and left bones for all paired bones (dentary, maxilla, premaxilla and palatine). For visualization of relevant data, see Online Resource 6.*

| Bones | Morph | $r^2$ | Slope | Intercept | p-value |
|---|---|---|---|---|---|
| Dentary | All | 0.709 | 0.826 | 2.44 | < 0.001 |
| | PL | 0.396 | 0.604 | 5.80 | < 0.001 |
| | PI | 0.571 | 0.771 | 3.94 | < 0.001 |
| | LB | 0.765 | 0.833 | 2.21 | < 0.001 |

| | | | | | |
|---|---|---|---|---|---|
| | SB | 0.292 | 0.544 | 5.85 | < 0.001 |
| Maxilla | All | 0.603 | 0.752 | 4.89 | < 0.001 |
| | PL | 0.397 | 0.675 | 6.12 | < 0.001 |
| | PI | 0.413 | 0.514 | 10.40 | < 0.001 |
| | LB | 0.467 | 0.834 | 3.70 | < 0.001 |
| | SB | 0.562 | 0.628 | 6.86 | < 0.001 |
| Premaxilla | All | 0.060 | 0.268 | 5.37 | < 0.001 |
| | PL | 0.074 | 0.253 | 5.48 | 0.054 |
| | PI | 0.081 | 0.295 | 5.39 | 0.053 |
| | LB | 0.046 | 0.213 | 5.55 | 0.147 |
| | SB | 0.044 | 0.244 | 5.54 | 0.087 |
| Palatine | All | 0.413 | 0.659 | 5.47 | < 0.001 |
| | PL | 0.278 | 0.580 | 6.35 | < 0.001 |
| | PI | 0.295 | 0.492 | 9.16 | < 0.001 |
| | LB | 0.478 | 0.802 | 2.95 | < 0.001 |
| | SB | 0.347 | 0.583 | 6.67 | < 0.001 |

***Online Resource 8*** *Results from student t-test, testing whether the mean asymmetry in tooth numbers differs from zero. Bonferroni corrected threshold (α = 0.01) to correct for tests on four bones.*

| | t | df | Estimated mean | p-value | Estimated skewness | Proportion of individuals with zero distribution |
|---|---|---|---|---|---|---|
| Dentary | -1.2763 | 237 | -0.1113445 | 0.2031 | -0.08090268 | 14.7% |
| Maxilla | -0.93919 | 227 | -0.1030702 | 0.3486 | -0.009834479 | 14.9% |
| Premaxilla | -2.2333 | 211 | -0.1533019 | 0.02658 | 0.008670065 | 21.2% |
| Palatine | -2.1043 | 224 | -0.2377778 | 0.03647 | -0.2141349 | 17.8% |

***Online Resource 9*** *Test of the influence of size (ln fork length), morph and the interaction on fluctuating asymmetry in tooth numbers in the four symmetric bones. Results from GLM with ln FL for size, the morphs (four types) and the interaction term.*

| Terms | | Dentary | Maxilla | Palatine | Premaxilla |
|---|---|---|---|---|---|
| FL effects (ln FL) | Deviance | 1.316 | 4.255 | 0.264 | 0.233 |
| | Resid. Df | 233 | 223 | 220 | 207 |
| | Resid. Dev | 421.55 | 615.40 | 621.06 | 208.31 |
| | p-value | 0.416 | 0.215 | 0.759 | 0.630 |
| | Deviance | 6.428 | 3.673 | 22.201 | 2.223 |

| | | | | | |
|---|---|---|---|---|---|
| Morph effects | Resid. Df | 234 | 224 | 221 | 208 |
| | Resid. Dev | 422.87 | 619.65 | 621.33 | 208.54 |
| | p-value | 0.308 | 0.723 | 0.048 | 0.530 |
| Morph x ln FL Interaction effect | Deviance | 10.644 | 5.805 | 11.875 | 2.979 |
| | Resid. Df | 230 | 220 | 217 | 204 |
| | Resid. Dev | 410.91 | 609.59 | 609.19 | 205.33 |
| | p-value | 0.114 | 0.553 | 0.238 | 0.398 |

P value, Bonferroni corrected threshold ($\alpha = 0.01$) to correct for tests on four bones.

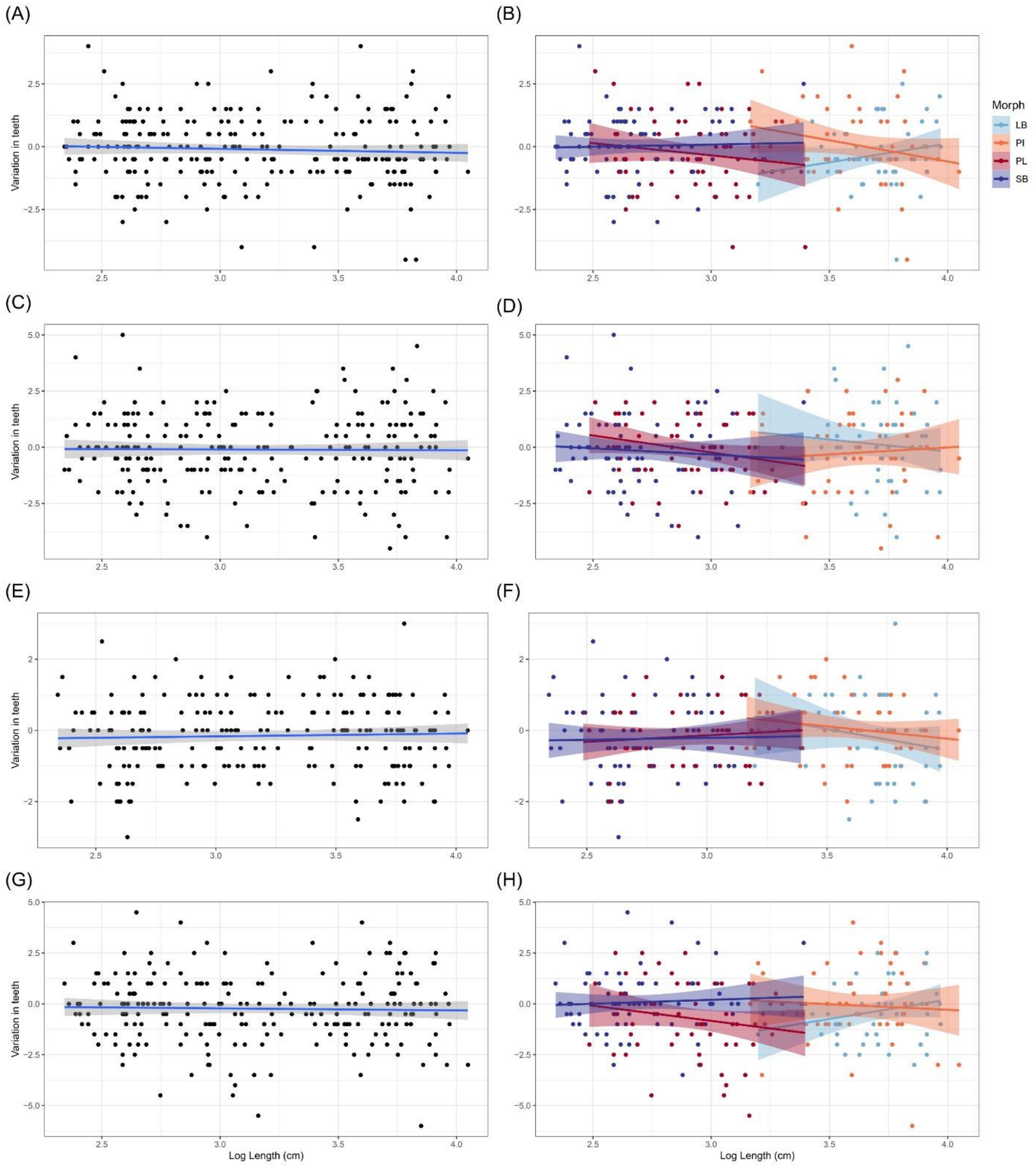


***Online Resource 10*** *The scatterplots show deviation in tooth numbers between left and right (Y-axis) against size (ln FL in cm, X-axis) for each of the four paired bones, dentary (A and B), maxilla (C and D), premaxilla (E and F) and palatine (G and H). The lines are a fitted linear model, with confidence intervals, nonsignificant by ANOVA on GLM models. Graphs on right (A, C, E and G) show all morph together and graphs on left (B, D, F and H) are coloured by morph.*

***Online Resource 11*** *Correlations (Pearson) between mean asymmetry in tooth numbers for each bone to each other and fork length.*

| | | Corr | p-value |
|---|---|---|---|
| Length | Dentary | - 0.052 | 0.9737 |
| | Maxilla | 0.002 | 0.4219 |
| | Premaxilla | 0.015 | 0.8237 |
| | Palatine | - 0.015 | 0.8282 |
| Dentary | Maxilla | 0.083 | 0.2134 |
| | Premaxilla | 0.037 | 0.5941 |
| | Palatine | -0.058 | 0.7562 |
| Maxilla | Premaxilla | 0.015 | 0.8309 |
| | Palatine | 0.021 | 0.7562 |
| Premaxilla | Palatine | 0.043 | 0.5509 |

***Online Resource 12*** *Average number of teeth for six bones and respective standard deviation (SD), calculated for all Arctic charr morphs. The numbers represent raw counts.*

| Morph | Sex | Dentary | Maxilla | Premaxilla | Palatine | Vomer | Glossohyal |
|---|---|---|---|---|---|---|---|
| PL | F | 15.68 ± 1.45 | 19.08 ± 1.63 | 7.60 ± 0.60 | 17.05 ± 1.71 | 4.70 ± 1.16 | 13.02 ± 1.22 |
| | M | 14.88 ± 1.46 | 18.97 ± 1.50 | 7.39 ± 0.66 | 16.11 ± 1.61 | 4.02 ± 1.30 | 12.17 ± 0.95 |
| | All | 15.18 ± 1.50 | 19.01 ± 1.54 | 7.47 ± 0.64 | 16.46 ± 1.70 | 4.27 ± 1.28 | 12.49 ± 1.13 |
| PI | F | 16.73 ± 1.44 | 21.84 ± 2.24 | 7.62 ± 0.41 | 18.41 ± 1.52 | 5.54 ± 1.31 | 12.68 ± 1.84 |
| | M | 17.06 ± 2.31 | 21.84 ± 1.76 | 7.60 ± 0.65 | 18.02 ± 1.82 | 5.83 ± 1.33 | 13.21 ± 2.14 |
| | All | 16.96 ± 2.08 | 21.84 ± 1.89 | 7.61 ± 0.59 | 18.14 ± 1.73 | 5.74 ± 1.31 | 13.05 ± 2.05 |
| LB | F | 15.35 ± 1.89 | 22.26 ± 1.76 | 7.65 ± 0.87 | 16.99 ± 1.52 | 4.32 ± 1.53 | 11.71 ± 1.13 |
| | M | 14.73 ± 2.59 | 21.08 ± 2.20 | 7.05 ± 0.71 | 15.93 ± 1.92 | 4.91 ± 1.23 | 11.58 ± 1.82 |
| | All | 14.94 ± 2.37 | 21.49 ± 2.12 | 7.26 ± 0.81 | 16.30 ± 1.85 | 4.70 ± 1.35 | 11.62 ± 1.60 |
| SB | F | 12.88 ± 1.08 | 18.64 ± 2.07 | 7.39 ± 0.69 | 15.58 ± 1.60 | 3.91 ± 1.35 | 11.32 ± 1.16 |
| | M | 12.70 ± 1.15 | 18.91 ± 2.02 | 7.50 ± 0.71 | 15.88 ± 1.17 | 3.85 ± 1.38 | 11.17 ± 1.53 |
| | All | 12.79 ± 1.10 | 18.79 ± 2.08 | 7.53 ± 0.72 | 15.74 ± 1.45 | 3.89 ± 1.33 | 11.27 ± 1.30 |
| All | F | 14.47 ± 2.05 | 19.79 ± 2.46 | 7.63 ± 0.69 | 16.55 ± 1.87 | 4.38 ± 1.44 | 11.97 ± 1.46 |
| | M | 14.84 ± 2.46 | 20.17 ± 2.26 | 7.36 ± 0.70 | 16.49 ± 1.86 | 4.67 ± 1.52 | 12.06 ± 1.80 |
| | All | 14.66 ± 2.30 | 20.01 ± 2.36 | 7.48 ± 0.70 | 16.52 ± 1.86 | 4.54 ± 1.48 | 12.02 ± 1.65 |

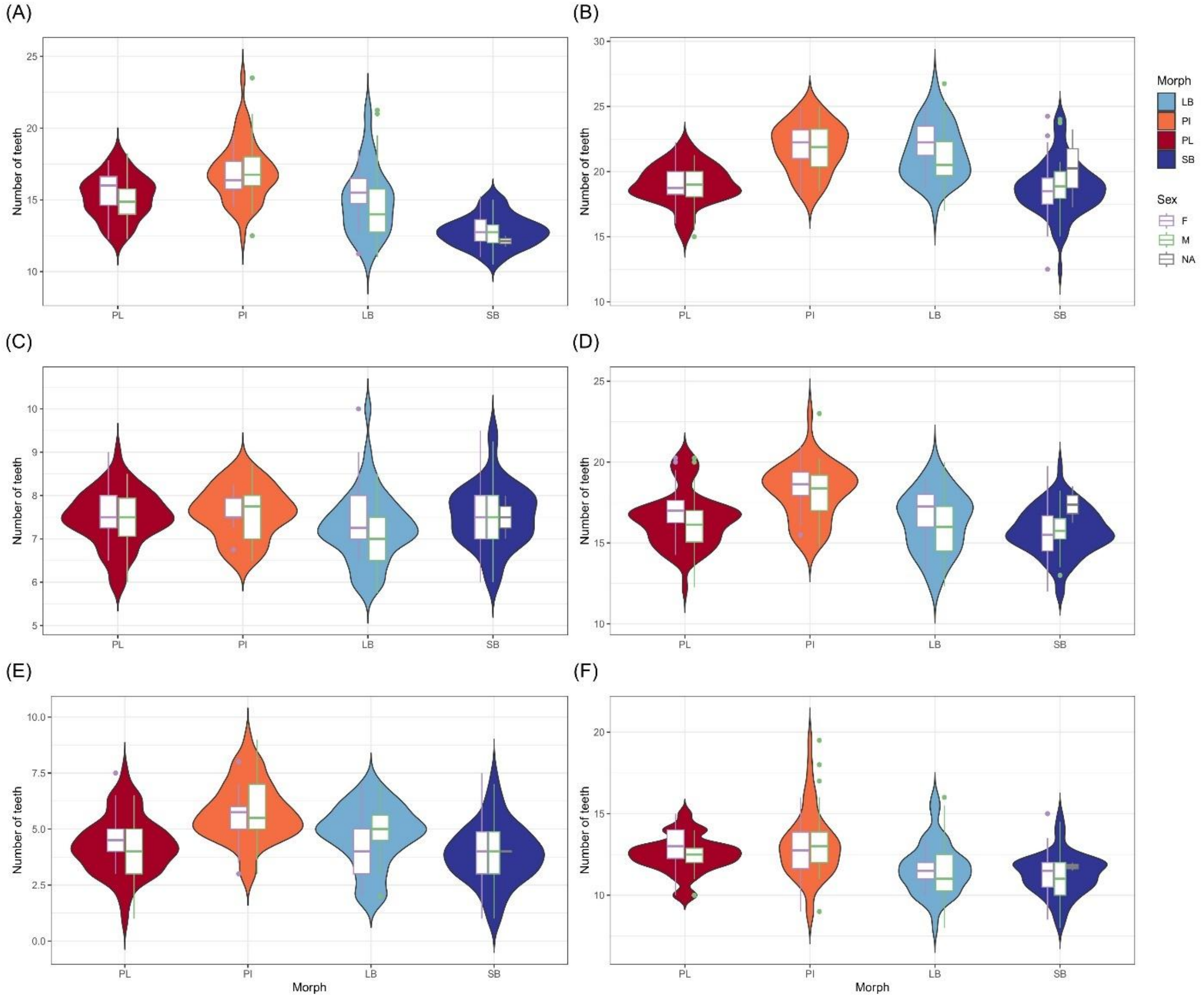


***Online Resource 13*** *Distribution of tooth number in six bones by morph. Also shown are sex specific distributions as boxplots. The Y-axis indicates the number of teeth (average per individual for the paired bones A-D, average over repeated measures for the non-paired E-F). The bones are: dentary (A), maxilla (B), premaxilla (C), palatine (D), vomer (E) and glossohyal bone (F). Morphs are coloured: SB: dark blue, LB: light blue, PI: orange, PL: red (see also inset). Superimposed are boxplots for the sexes (female: purple, males: green, not sexed: grey, only in SB-charr).*

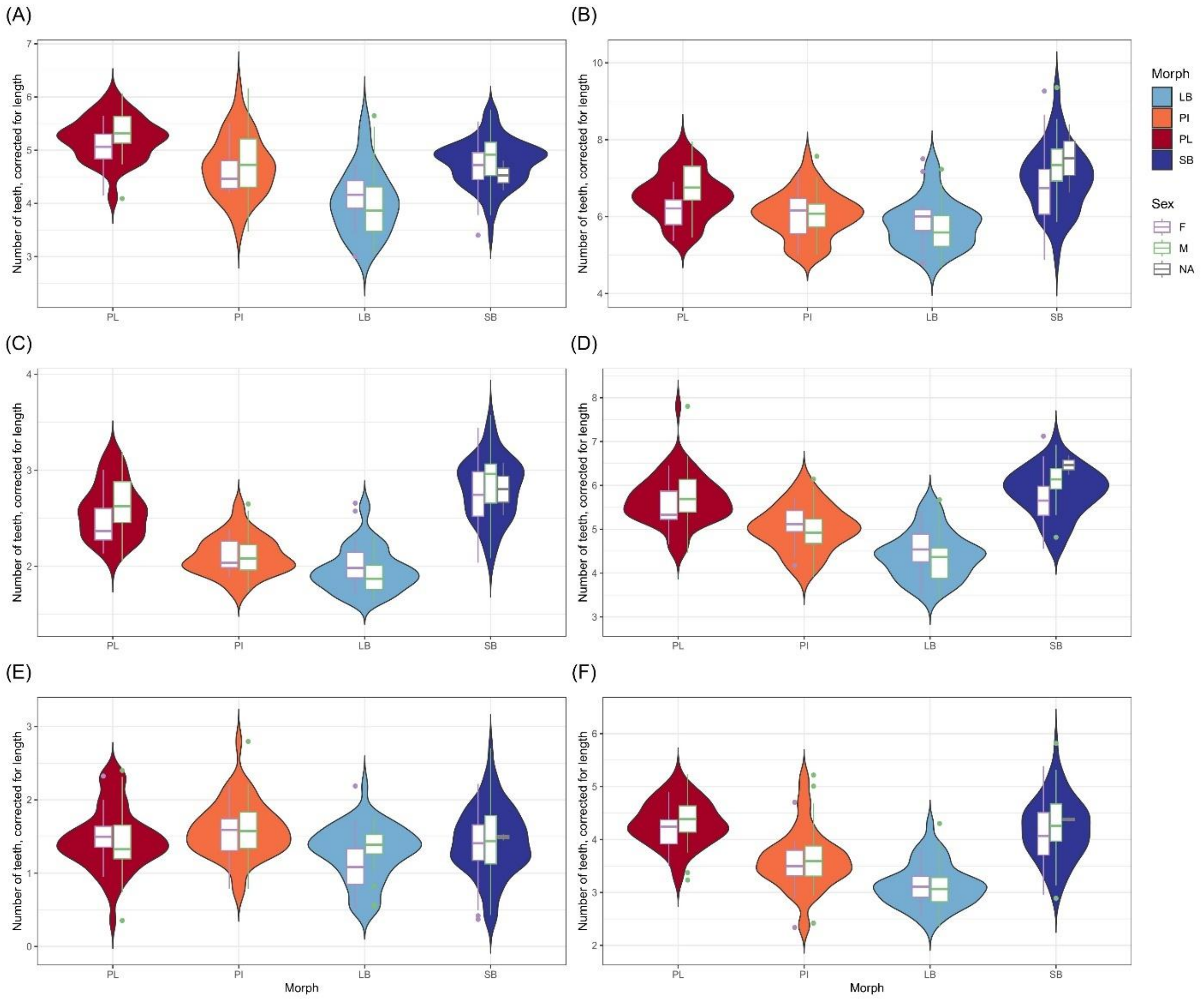


***Online Resource 14*** *The size corrected distribution of tooth number in six bones by morph. Also shown are sex specific distributions as boxplots. The Y-axis indicates the number of teeth corrected for with individual length (FL) (teeth/length). The bones are: dentary (A), maxilla (B), premaxilla (C), palatine (D), vomer (E) and glossohyal bone (F). Morph are coloured: SB: dark blue, LB: light blue, PI: orange, PL: red (see also inset). Superimposed are boxplots for the sexes (females: purple, males: green, not sexed: grey, only in SB-charr).*

***Online Resource 15*** *Result from emtrends, showing the different trends between tooth number and different length measurements (raw fork length, ln fork length, ln C-size).*

| Bone | Length measurement | trends | se | df | Lower.CI | Upper.CI |
|---|---|---|---|---|---|---|
| Dentary | Raw fork length | 0.122 | 0.015 | 4564 | 0.093 | 0.152 |
| Maxilla | Raw fork length | 0.069 | 0.015 | 4564 | 0.039 | 0.099 |
| Premaxilla | Raw fork length | 0.031 | 0.015 | 4564 | 0.001 | 0.061 |
| Palatine | Raw fork length | 0.078 | 0.015 | 4564 | 0.048 | 0.108 |
| Vomer | Raw fork length | 0.060 | 0.018 | 4564 | 0.025 | 0.094 |
| Glossohyal | Raw fork length | 0.063 | 0.018 | 4564 | 0.028 | 0.097 |

| | | | | | | |
|---|---|---|---|---|---|---|
| Dentary | ln fork length | 2.010 | 0.269 | 4564 | 1.482 | 2.537 |
| Maxilla | ln fork length | 1.087 | 0.270 | 4564 | 0.557 | 1.617 |
| Premaxilla | ln fork length | 0.223 | 0.271 | 4564 | -0.308 | 0.755 |
| Palatine | ln fork length | 1.375 | 0.271 | 4564 | 0.844 | 1.906 |
| Vomer | ln fork length | 1.104 | 0.321 | 4564 | 0.475 | 1.733 |
| Glossohyal | ln fork length | 0.803 | 0.315 | 4564 | 0.184 | 1.421 |
| Dentary | ln C-size dentary | 3.327 | 0.325 | 4548 | 2.691 | 3.960 |
| Maxilla | ln C-size maxilla | 2.286 | 0.330 | 4548 | 1.639 | 2.930 |
| Premaxilla | ln C-size premaxilla | 0.615 | 0.298 | 4564 | 0.031 | 1.200 |

***Online Resource 16*** *The influence of size, morph, sex and side (left and right) on tooth number in six jaw bones. Analysed with generalized liner mixed model (glmmTMB) using ln C-size for size.*

| Terms | | Dentary | Maxilla | Premaxilla |
|---|---|---|---|---|
| Morph Effect | $\chi^2$ | 88.505 | 3.614 | 16.237 |
| | df | 3 | 3 | 3 |
| | p-value | **< 0.001** | 0.306 | **0.001** |
| Sex Effect | $\chi^2$ | 0.666 | 0.001 | 6.862 |
| | df | 1 | 1 | 1 |
| | p-value | 0.415 | 0.974 | **0.008** |
| Bone size | $\chi^2$ | 55.969 | 22.630 | 7.484 |
| | df | 1 | 1 | 1 |
| | p-value | **< 0.001** | **< 0.001** | **0.006** |
| Side | $\chi^2$ | 1.762 | 0.906 | 5.322 |
| | df | 1 | 1 | 1 |
| | p-value | 0.184 | 0.341 | 0.021 |
| Morph x Sex Interaction Effect | $\chi^2$ | 3.691 | 8.330 | 4.118 |
| | df | 3 | 3 | 3 |
| | p-value | 0.297 | 0.040 | 0.249 |
| Morph x size Interaction Effect | $\chi^2$ | 3.473 | 3.183 | 3.213 |
| | df | 3 | 3 | 3 |
| | P-value | 0.3243 | 0.364 | 0.360 |
| Sex x Size Interaction Effect | $\chi^2$ | 0.255 | 1.805 | 0.270 |
| | df | 1 | 1 | 1 |
| | p-value | 0.614 | 0.179 | 0.604 |

P value, Bonferroni corrected threshold, $\alpha = 0.008$

***Online Resource 17*** *Pairwise comparisons of teeth numbers between morphs, results from the generalized liner mixed models (glmmTMB), when using ln FL for size.*

| Bones | Morph pairs | Estimates | se | lower. CI | upper. CI | t.ratio | p-value |
|---|---|---|---|---|---|---|---|
| Dentary | LB-PI | -2.360 | 0.82 | -4.476 | -0.237 | -2.86 | 0.022 |
| | PL-SB | 3.100 | 0.68 | 1.349 | 4.843 | 4.56 | **< 0.001** |
| | LB-PL | -3.370 | 0.79 | -5.402 | -1.331 | -4.26 | **< 0.001** |
| | PI-SB | 2.090 | 0.68 | 0.336 | 3.836 | 3.07 | 0.989 |
| | LB-SB | -0.270 | 0.84 | -2.441 | 1.900 | -0.32 | 0.989 |
| | PI-PL | -1.010 | 0.65 | -2.680 | 0.660 | -1.56 | 0.404 |
| Maxilla | LB-PI | 0.051 | 0.93 | -2.354 | 2.460 | 0.05 | 1.000 |
| | PL-SB | -0.278 | 0.80 | -2.332 | 1.780 | -0.35 | 0.986 |
| | LB-PL | 1.652 | 0.91 | -0.698 | 4.000 | 1.81 | 0.270 |
| | PI-SB | 1.324 | 0.79 | -0.705 | 3.350 | 1.68 | 0.335 |
| | LB-SB | 1.375 | 0.97 | -1.129 | 3.880 | 1.41 | 0.491 |
| | PI-PL | 1.602 | 0.75 | -0.321 | 3.520 | 2.14 | 0.140 |
| Palatine | LB-PI | -1.252 | 0.77 | -3.240 | 0.732 | -1.62 | 0.366 |
| | PL-SB | 0.398 | 0.68 | -1.360 | 2.153 | 0.58 | 0.937 |
| | LB-PL | -1.251 | 0.76 | -3.200 | 0.699 | -1.65 | 0.350 |
| | PI-SB | 0.398 | 0.66 | -1.310 | 2.103 | 0.60 | 0.932 |
| | LB-SB | -0.854 | 0.81 | -2.950 | 1.240 | -1.05 | 0.720 |
| | PI-PL | 0.000 | 0.63 | -1.610 | 1.613 | 0.00 | 1.000 |
| Premaxilla | LB-PI | -0.356 | 0.31 | -1.162 | 0.451 | -1.14 | 0.668 |
| | PL-SB | -0.156 | 0.29 | -0.889 | 0.576 | -0.55 | 0.947 |
| | LB-PL | -0.372 | 0.31 | -1.176 | 0.432 | -1.19 | 0.633 |
| | PI-SB | -0.173 | 0.27 | -0.876 | 0.531 | -0.63 | 0.922 |
| | LB-SB | -0.529 | 0.33 | -1.386 | 0.330 | -1.59 | 0.387 |
| | PI-PL | -0.016 | 0.26 | -0.683 | 0.650 | -0.06 | 1.000 |
| Vomer | LB-PI | -1.108 | 0.63 | -2.734 | 0.518 | -1.76 | 0.295 |
| | PL-SB | -0.274 | 0.59 | -1.792 | 1.245 | -0.47 | 0.967 |
| | LB-PL | -0.345 | 0.64 | -1.990 | 1.301 | -0.54 | 0.949 |
| | PI-SB | 0.490 | 0.55 | -0.936 | 1.915 | 0.89 | 0.812 |
| | LB-SB | -0.618 | 0.67 | -2.337 | 1.101 | -0.93 | 0.790 |
| | PI-PL | 0.763 | 0.54 | -0.636 | 2.163 | 1.41 | 0.496 |
| Glossohyal | LB-PI | -2.286 | 0.75 | -4.208 | -0.363 | -3.07 | 0.012 |
| | PL-SB | 1.378 | 0.63 | -0.258 | 3.014 | 2.17 | 0.133 |
| | LB-PL | -2.507 | 0.73 | -4.381 | -0.634 | -3.45 | **0.003** |
| | PI-SB | 1.156 | 0.63 | -0.458 | 2.771 | 1.85 | 0.253 |
| | LB-SB | -1.130 | 0.78 | -3.129 | 0.870 | -1.46 | 0.465 |
| | PI-PL | -0.222 | 0.59 | -1.753 | 1.309 | -0.37 | 0.982 |

***Online Resource 18*** *Pairwise comparisons of teeth numbers between morphs within sex, results from the generalized liner mixed models (glmmTMB), when using ln FL for size.*

| Bones | Sex | Morph pairs | Estimates | se | lower. CI | upper. CI | t.ratio | p-value |
|---|---|---|---|---|---|---|---|---|
| Dentary | Females | LB-PI | -1.923 | 0.96 | -4.386 | 0.539 | -2.01 | 0.185 |
| | | PL-SB | 2.721 | 0.60 | 1.188 | 4.254 | 4.57 | **< 0.001** |
| | | LB-PL | -2.841 | 0.97 | -5.328 | -0.354 | -2.94 | 0.018 |
| | | PI-SB | 1.803 | 0.92 | -0.552 | 4.158 | 1.97 | 0.200 |
| | | LB-SB | -0.120 | 1.01 | -2.729 | 2.489 | -0.12 | 0.999 |
| | | PI-PL | -0.918 | 0.86 | -3.141 | 1.306 | -1.06 | 0.713 |
| | Males | LB-PI | -2.789 | 0.83 | -4.922 | -0.656 | -3.37 | **0.004** |
| | | PL-SB | 3.471 | 0.94 | 1.046 | 5.897 | 3.68 | **0.001** |
| | | LB-PL | -3.892 | 0.96 | -6.371 | -1.412 | -4.04 | **< 0.001** |
| | | PI-SB | 2.369 | 0.99 | -0.170 | 4.908 | 2.40 | 0.078 |
| | | LB-SB | -0.420 | 1.18 | -3.457 | 2.616 | -0.36 | 0.985 |
| | | PI-PL | -1.102 | 0.80 | -3.161 | 0.956 | -1.38 | 0.513 |
| Maxilla | Females | LB-PI | 0.717 | 1.09 | -2.089 | 3.520 | 0.66 | 0.913 |
| | | PL-SB | -0.073 | 0.70 | -1.871 | 1.730 | -0.10 | 1.000 |
| | | LB-PL | 3.124 | 1.11 | 0.268 | 5.980 | 2.82 | 0.026 |
| | | PI-SB | 2.335 | 1.06 | -0.387 | 5.030 | 2.21 | 0.122 |
| | | LB-SB | 3.052 | 1.16 | 0.059 | 6.040 | 2.63 | 0.044 |
| | | PI-PL | 2.407 | 2.41 | -0.159 | 4.970 | 2.41 | 0.075 |
| | Males | LB-PI | -0.615 | 0.94 | -3.038 | 1.810 | -0.65 | 0.914 |
| | | PL-SB | -0.483 | 1.10 | -3.322 | 2.360 | -0.44 | 0.972 |
| | | LB-PL | 0.180 | 1.12 | -2.689 | 3.050 | 0.16 | 0.999 |
| | | PI-SB | 0.313 | 1.14 | -2.628 | 3.250 | 0.27 | 0.993 |
| | | LB-SB | -0.303 | 1.37 | -3.822 | 3.220 | -0.22 | 0.996 |
| | | PI-PL | 0.796 | 0.92 | -1.565 | 3.160 | 0.87 | 0.822 |
| Palatine | Females | LB-PI | -0.935 | 0.90 | -3.254 | 1.384 | -1.04 | 0.727 |
| | | PL-SB | 0.651 | 0.60 | -0.885 | 2.188 | 1.09 | 0.695 |
| | | LB-PL | -0.668 | 0.92 | -3.044 | 1.709 | -0.72 | 0.888 |
| | | PI-SB | 0.919 | 0.89 | -1.363 | 3.201 | 1.04 | 0.728 |
| | | LB-SB | -0.016 | 0.97 | -2.519 | 2.487 | -0.02 | 1.000 |
| | | PI-PL | 0.268 | 0.83 | -1.875 | 2.410 | 0.32 | 0.989 |
| | Males | LB-PI | -1.568 | 0.78 | -3.572 | 0.436 | -2.01 | 0.183 |
| | | PL-SB | 0.144 | 0.94 | -2.271 | 2.560 | 0.15 | 0.999 |
| | | LB-PL | -1.835 | 0.93 | -4.227 | 0.557 | -1.98 | 0.198 |
| | | PI-SB | -0.122 | 0.96 | -2.593 | 2.348 | -0.13 | 0.999 |
| | | LB-SB | -1.691 | 1.15 | -4.639 | 1.258 | -1.48 | 0.453 |
| | | PI-PL | -0.267 | 0.78 | -2.260 | 1.727 | -0.34 | 0.986 |
| Premaxilla | Females | LB-PI | -0.087 | 0.37 | -1.036 | 0.863 | -0.24 | 0.995 |

| | | | | | | | | |
|---|---|---|---|---|---|---|---|---|
| | | PL-SB | -0.221 | 0.25 | -0.858 | 0.416 | -0.89 | 0.809 |
| | | LB-PL | -0.292 | 0.38 | -1.272 | 0.688 | -0.77 | 0.870 |
| | | PI-SB | -0.426 | 0.36 | -1.361 | 0.508 | -1.17 | 0.643 |
| | | LB-SB | -0.513 | 0.40 | -1.536 | 0.511 | -1.29 | 0.570 |
| | | PI-PL | -0.205 | 0.34 | -1.091 | 0.681 | -0.60 | 0.933 |
| | Males | LB-PI | -0.625 | 0.32 | -1.439 | 0.189 | -1.98 | 0.198 |
| | | PL-SB | -0.092 | 0.39 | -1.104 | 0.921 | -0.23 | 0.996 |
| | | LB-PL | -0.452 | 0.39 | -1.444 | 0.539 | -1.18 | 0.643 |
| | | PI-SB | 0.081 | 0.40 | -0.944 | 1.106 | 0.20 | 0.997 |
| | | LB-SB | -0.544 | 0.47 | -1.762 | 0.673 | -1.15 | 0.658 |
| | | PI-PL | 0.173 | 0.32 | -0.656 | 1.001 | 0.54 | 0.950 |
| Vomer | Females | LB-PI | -1.254 | 0.74 | -3.160 | 0.648 | -1.70 | 0.325 |
| | | PL-SB | -0.027 | 0.51 | -1.340 | 1.290 | -0.05 | 1.000 |
| | | LB-PL | -0.756 | 0.78 | -2.760 | 1.247 | -0.97 | 0.765 |
| | | PI-SB | 0.472 | 0.73 | -1.410 | 2.354 | 0.65 | 0.917 |
| | | LB-SB | -0.783 | 0.80 | -2.840 | 1.270 | -0.98 | 0.759 |
| | | PI-PL | 0.499 | 0.71 | -1.330 | 2.325 | 0.70 | 0.896 |
| | Males | LB-PI | -0.962 | 0.64 | -2.610 | 0.685 | -1.51 | 0.434 |
| | | PL-SB | -0.520 | 0.81 | -2.600 | 1.564 | -0.64 | 0.918 |
| | | LB-PL | 0.066 | 0.79 | -1.970 | 2.102 | 0.08 | 1.000 |
| | | PI-SB | 0.508 | 0.81 | -1.590 | 2.608 | 0.62 | 0.924 |
| | | LB-SB | -0.454 | 0.96 | -2.920 | 2.010 | -0.48 | 0.965 |
| | | PI-PL | 1.028 | 0.67 | -0.710 | 2.767 | 1.53 | 0.423 |
| Glossohyal | Females | LB-PI | -1.938 | 0.87 | -4.172 | 0.297 | -2.24 | 0.115 |
| | | PL-SB | 1.836 | 0.55 | 0.407 | 3.265 | 3.31 | **0.006** |
| | | LB-PL | -2.304 | 0.88 | -4.579 | -0.028 | -2.61 | 0.046 |
| | | PI-SB | 1.470 | 0.84 | -0.692 | 3.632 | 1.75 | 0.298 |
| | | LB-SB | -0.468 | 0.92 | -2.844 | 1.909 | -0.51 | 0.957 |
| | | PI-PL | -0.366 | 0.79 | -2.414 | 1.682 | -0.46 | 0.968 |
| | Males | LB-PI | -2.634 | 0.75 | -4.579 | -0.690 | -3.49 | **0.003** |
| | | PL-SB | 0.920 | 0.88 | -1.342 | 3.182 | 1.05 | 0.721 |
| | | LB-PL | -2.711 | 0.89 | -5.008 | -0.415 | -3.05 | 0.013 |
| | | PI-SB | 0.843 | 0.91 | -1.503 | 3.188 | 0.93 | 0.791 |
| | | LB-SB | -1.791 | 1.09 | -4.614 | 1.031 | -1.64 | 0.359 |
| | | PI-PL | -0.077 | 0.73 | -1.957 | 1.803 | -0.11 | 1.000 |

***Online Resource 19*** *Pairwise comparisons of teeth numbers between sexes within morph, results from the generalized liner mixed models (glmmTMB), when using ln FL for size.*

| Bones | Morph | Pairs | Estimates | se | lower. CI | upper. CI | t.ratio | p-value |
|---|---|---|---|---|---|---|---|---|
| Dentary | PL | F-M | -0.685 | 0.62 | -1.900 | 0.533 | -1.10 | 0.270 |
| | PI | F-M | -0.500 | 0.67 | -1.810 | 0.811 | -0.75 | 0.454 |
| | LB | F-M | 0.366 | 0.72 | -1.050 | 1.781 | 0.51 | 0.612 |
| | SB | F-M | 0.065 | 0.83 | -1.560 | 1.687 | 0.08 | 0.937 |
| Maxilla | PL | F-M | -1.125 | 0.72 | -2.535 | 0.286 | -1.57 | 0.118 |
| | PI | F-M | 0.487 | 0.76 | -1.013 | 1.987 | 0.64 | 0.524 |
| | LB | F-M | 1.820 | 0.83 | 0.199 | 3.440 | 2.20 | 0.028 |
| | SB | F-M | -1.535 | 0.96 | -3.426 | 0.357 | -1.59 | 0.112 |
| Palatine | PL | F-M | -0.174 | 0.61 | -1.369 | 1.022 | -0.29 | 0.776 |
| | PI | F-M | 0.361 | 0.64 | -0.887 | 1.608 | 0.57 | 0.571 |
| | LB | F-M | 0.994 | 0.69 | -0.351 | 2.340 | 1.45 | 0.147 |
| | SB | F-M | -0.680 | 0.82 | -2.280 | 0.919 | -0.84 | 0.404 |
| Premaxilla | PL | F-M | 0.250 | 0.26 | -0.256 | 0.755 | 0.97 | 0.333 |
| | PI | F-M | -0.128 | 0.26 | -0.640 | 0.383 | -0.49 | 0.623 |
| | LB | F-M | 0.410 | 0.28 | -0.146 | 0.966 | 1.45 | 0.148 |
| | SB | F-M | 0.379 | 0.34 | -0.286 | 1.044 | 1.12 | 0.264 |
| Vomer | PL | F-M | 0.228 | 0.53 | -0.815 | 1.271 | 0.43 | 0.668 |
| | PI | F-M | -0.302 | 0.53 | -1.338 | 0.735 | -0.57 | 0.568 |
| | LB | F-M | -0.593 | 0.57 | -1.713 | 0.526 | -1.04 | 0.298 |
| | SB | F-M | -0.265 | 0.69 | -1.617 | 1.087 | -0.39 | 0.700 |
| Glossohyal | PL | F-M | 0.226 | 0.57 | -0.898 | 1.350 | 0.40 | 0.693 |
| | PI | F-M | -0.062 | 0.61 | -1.258 | 1.134 | -0.10 | 0.918 |
| | LB | F-M | 0.634 | 0.66 | -0.664 | 1.932 | 0.96 | 0.338 |
| | SB | F-M | -0.690 | 0.77 | -2.194 | 0.815 | -0.90 | 0.368 |

***Online Resource 20*** *Pairwise comparisons of teeth numbers between morphs, results from the generalized liner mixed models (glmmTMB), when using ln C-size for size.*

| Bones | Morph pairs | Estimates | se | lower. CI | upper. CI | t.ratio | p-value |
|---|---|---|---|---|---|---|---|
| Dentary | LB-PI | -2.410 | 0.79 | -4.433 | -0.379 | -3.06 | 0.012 |
| | PL-SB | 2.350 | 0.52 | 1.002 | 3.691 | 4.49 | **< 0.001** |
| | LB-PL | -3.440 | 0.63 | -5.067 | -1.814 | -5.45 | **< 0.001** |
| | PI-SB | 1.310 | 0.68 | -0.436 | 3.061 | 1.93 | 0.216 |
| | LB-SB | -1.090 | 0.69 | -2.862 | 0.673 | -1.59 | 0.383 |
| | PI-PL | -1.030 | 0.65 | -2.708 | 0.639 | -1.59 | 0.384 |
| Maxilla | LB-PI | -0.939 | 0.92 | -3.295 | 1.420 | -1.03 | 0.734 |
| | PL-SB | 0.082 | 0.62 | -1.505 | 1.670 | 0.13 | 0.999 |

| | | | | | | | |
|---|---|---|---|---|---|---|---|
| | LB-PL | 0.247 | 0.80 | -1.813 | 2.310 | 0.31 | 0.990 |
| | PI-SB | 1.267 | 0.74 | -0.637 | 3.170 | 1.71 | 0.318 |
| | LB-SB | 0.328 | 0.82 | -1.776 | 2.430 | 0.40 | 0.978 |
| | PI-PL | 1.185 | 0.74 | -0.731 | 3.100 | 1.59 | 0.384 |
| Premaxilla | LB-PI | -0.672 | 0.30 | -1.438 | 0.094 | -2.26 | 0.109 |
| | PL-SB | -0.211 | 0.23 | -0.801 | 0.380 | -0.92 | 0.795 |
| | LB-PL | -0.737 | 0.29 | -1.494 | 0.021 | -2.50 | 0.060 |
| | PI-SB | -0.276 | 0.23 | -0.864 | 0.313 | -1.21 | 0.624 |
| | LB-SB | -0.947 | 0.28 | -1.664 | -0.230 | -3.40 | **0.004** |
| | PI-PL | -0.065 | 0.25 | -0.713 | 0.583 | -0.26 | 0.994 |

***Online Resource 21*** *Pairwise comparisons of teeth numbers between morphs within sex, results from the generalized liner mixed models (glmmTMB), when using ln C-size for size.*

| Bones | Sex | Morph pairs | Estimates | se | lower. CI | upper. CI | t.ratio | p-value |
|---|---|---|---|---|---|---|---|---|
| Dentary | Females | LB-PI | -1.892 | 0.85 | -4.080 | 0.295 | -2.23 | 0.117 |
| | | PL-SB | 2.395 | 0.53 | 1.032 | 3.758 | 4.52 | **< 0.001** |
| | | LB-PL | -2.686 | 0.78 | -4.682 | -0.691 | -3.47 | **0.003** |
| | | PI-SB | 1.601 | 0.92 | -0.772 | 3.974 | 1.74 | 0.306 |
| | | LB-SB | -0.292 | 0.83 | -2.421 | 1.837 | -0.35 | 0.985 |
| | | PI-PL | -0.794 | 0.87 | -3.031 | 1.443 | -0.91 | 0.798 |
| | Males | LB-PI | -2.920 | 0.87 | -5.148 | -0.692 | -3.37 | **0.004** |
| | | PL-SB | 2.298 | 0.69 | 0.528 | 4.068 | 3.34 | **0.005** |
| | | LB-PL | -4.195 | 0.81 | -6.285 | -2.104 | -5.16 | **< 0.001** |
| | | PI-SB | 1.024 | 0.89 | -1.266 | 3.313 | 1.15 | 0.658 |
| | | LB-SB | -1.896 | 0.97 | -4.380 | 0.587 | -1.97 | 0.202 |
| | | PI-PL | -1.275 | 0.77 | -3.252 | 0.703 | -1.66 | 0.346 |
| Maxilla | Females | LB-PI | -0.287 | 1.00 | -2.847 | 2.273 | -0.29 | 0.992 |
| | | PL-SB | 0.191 | 0.62 | -1.398 | 1.779 | 0.31 | 0.990 |
| | | LB-PL | 1.870 | 0.98 | -0.642 | 4.383 | 1.92 | 0.222 |
| | | PI-SB | 2.348 | 1.02 | -0.282 | 4.978 | 2.30 | 0.099 |
| | | LB-SB | 2.061 | 0.99 | -0.486 | 4.608 | 2.08 | 0.160 |
| | | PI-PL | 2.158 | 1.01 | -0.430 | 4.745 | 2.15 | 0.139 |
| | Males | LB-PI | -1.591 | 1.00 | -4.163 | 0.981 | -1.59 | 0.384 |
| | | PL-SB | -0.027 | 0.80 | -2.090 | 2.035 | -0.03 | 1.000 |
| | | LB-PL | -1.377 | 1.01 | -3.964 | 1.209 | -1.37 | 0.518 |
| | | PI-SB | 0.186 | 0.96 | -2.290 | 2.662 | 0.19 | 0.997 |
| | | LB-SB | -1.405 | 1.13 | -4.308 | 1.498 | -1.25 | 0.598 |
| | | PI-PL | 0.213 | 0.87 | -2.023 | 2.449 | 0.25 | 0.995 |

| | | | | | | | | |
|---|---|---|---|---|---|---|---|---|
| Premaxilla | Females | LB-PI | -0.384 | 0.33 | -1.228 | 0.460 | -1.17 | 0.645 |
| | | PL-SB | -0.215 | 0.22 | -0.789 | 0.359 | -0.97 | 0.769 |
| | | LB-PL | -0.570 | 0.36 | -1.484 | 0.344 | -1.60 | 0.377 |
| | | PI-SB | -0.401 | 0.30 | -1.182 | 0.381 | -1.32 | 0.551 |
| | | LB-SB | -0.785 | 0.34 | -1.646 | 0.077 | -2.35 | 0.089 |
| | | PI-PL | -0.186 | 0.33 | -1.027 | 0.656 | -0.57 | 0.942 |
| | Males | LB-PI | -0.959 | 0.33 | -1.814 | -0.105 | -2.89 | 0.021 |
| | | PL-SB | -0.206 | 0.29 | -0.964 | 0.552 | -0.70 | 0.897 |
| | | LB-PL | -0.904 | 0.38 | -1.879 | 0.072 | -2.38 | 0.081 |
| | | PI-SB | -0.151 | 0.31 | -0.960 | 0.659 | -0.48 | 0.964 |
| | | LB-SB | -1.110 | 0.38 | -2.088 | -0.131 | -2.92 | 0.019 |
| | | PI-PL | 0.056 | 0.32 | -0.777 | 0.889 | 0.17 | 0.998 |

***Online Resource 22*** *Pairwise comparisons of teeth numbers between sexes within morph, results from the generalized liner mixed models (glmmTMB), when using ln C-size for size.*

| Bones | Morph | Pairs | Estimates | se | lower. CI | upper. CI | t.ratio | p-value |
|---|---|---|---|---|---|---|---|---|
| Dentary | PL | F-M | -0.213 | 0.50 | -1.192 | 0.765 | -0.43 | 0.669 |
| | PI | F-M | 0.267 | 0.73 | -1.159 | 1.694 | 0.37 | 0.713 |
| | LB | F-M | 1.295 | 0.68 | -0.047 | 2.637 | 1.89 | 0.059 |
| | SB | F-M | -0.310 | 0.61 | -1.512 | 0.892 | -0.51 | 0.613 |
| Maxilla | PL | F-M | -1.039 | 0.60 | -2.206 | 0.128 | -1.75 | 0.081 |
| | PI | F-M | 0.906 | 0.81 | -0.674 | 2.485 | 1.13 | 0.261 |
| | LB | F-M | 2.209 | 0.82 | 0.600 | 3.818 | 2.70 | **0.007** |
| | SB | F-M | -1.257 | 0.69 | -2.613 | 0.099 | -1.82 | 0.069 |
| Premaxilla | PL | F-M | 0.220 | 0.24 | -0.246 | 0.686 | 0.93 | 0.355 |
| | PI | F-M | -0.022 | 0.26 | -0.535 | 0.491 | -0.08 | 0.934 |
| | LB | F-M | 0.554 | 0.29 | -0.006 | 1.113 | 1.94 | 0.052 |
| | SB | F-M | 0.228 | 0.22 | -0.212 | 0.669 | 1.02 | 0.309 |

***Online Resource 23*** *Linear regression (function lm) examining the repeatability of the two methods (fronto/lateral method and medial/ventral method) used when measuring the angle of maxillary teeth. This was analysed on 62 specimens from 3 morphs. The Medial/ventral method appeared to be more repeatable.*

| Method | Side | $r^2$ | Estimate (Slope) | Estimate (Intersect) | t-value | p-value |
|---|---|---|---|---|---|---|
| Fronto/lateral | Left | 0.689 | 0.89 | 2.92 | 11.67 | < 0.001 |
| Fronto/lateral | Right | 0.480 | 0.69 | 14.50 | 7.58 | < 0.001 |
| Medial/ventral | Left | 0.716 | 0.80 | 8.38 | 12.45 | < 0.001 |
| Medial/ventral | Right | 0.730 | 0.78 | 8.44 | 12.89 | < 0.001 |

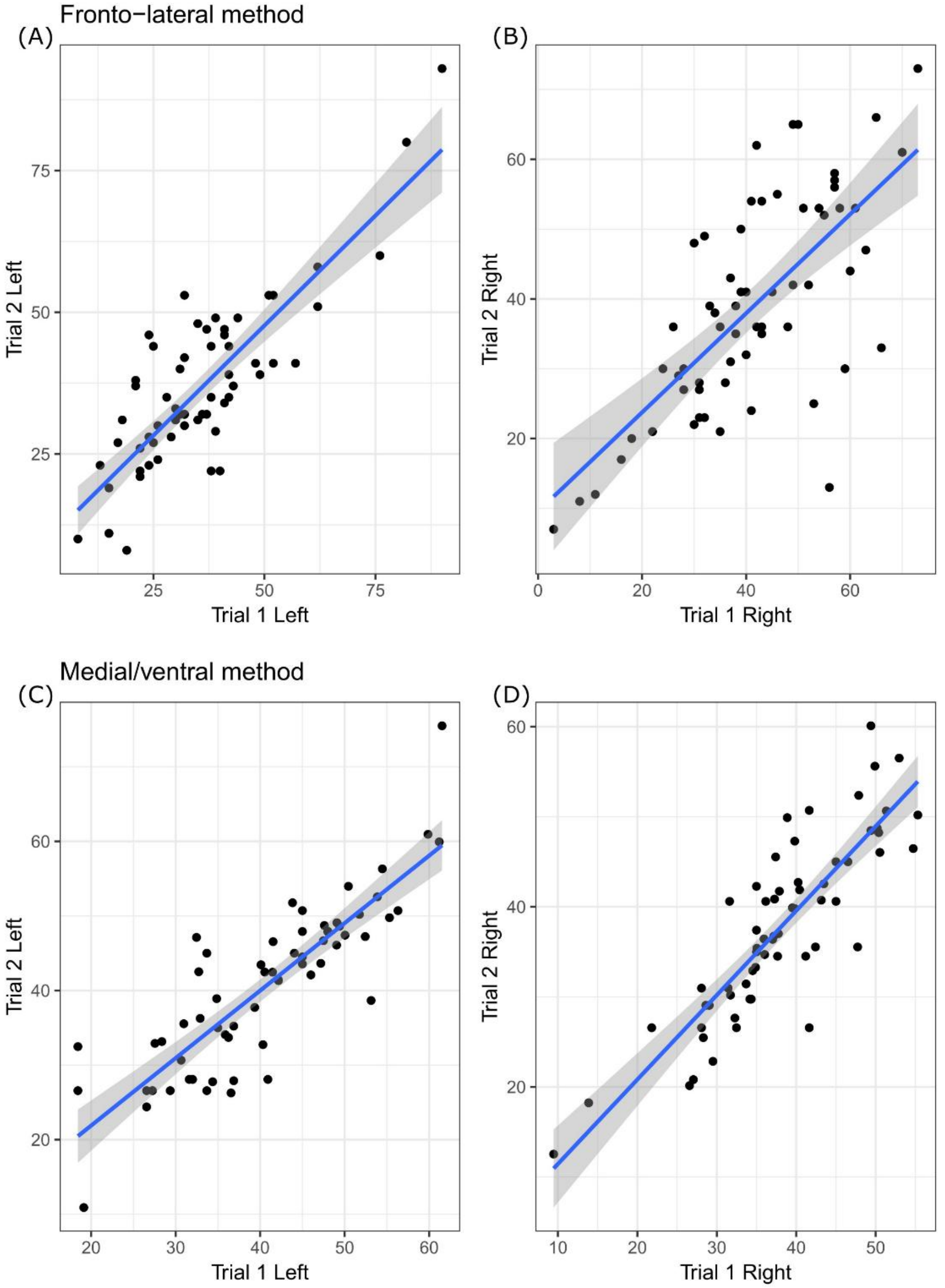


***Online Resource 24*** *Test of repeatability for estimation of maxillary teeth angle with linear regression. Scatterplots show the liner relationship between the two trials for both methods (fronto/lateral method and medial/ventral method) used to measure the angle. This was analysed on 62 specimens from 3 morphs. The Medial/ventral method appeared to be more repeatable.*

***Online Resource 25*** *Average angle of maxillary teeth (± standard deviation) for each morph, also by sex.*

| Morph | Sex | 1st tooth | 5th tooth | Mean |
|---|---|---|---|---|
| PL | F | 41.96 ± 7.74 | 36.27 ± 10.35 | 39.11 ± 7.75 |
| | M | 28.30 ± 9.10 | 27.72 ± 9.20 | 28.01 ± 8.59 |
| | All | 33.76 ± 10.86 | 31.14 ± 10.47 | 32.45 ± 9.86 |
| PI | F | 41.96 ± 7.74 | 45.80 ± 7.49 | 43.82 ± 6.55 |
| | M | 38.27 ± 7.34 | 40.67 ± 8.38 | 39.47 ± 6.77 |
| | All | 39.38 ± 7.47 | 42.27 ± 8.38 | 40.82 ± 6.94 |
| LB | F | 49.83 ± 6.82 | 50.83 ± 5.15 | 50.33 ± 5.52 |
| | M | 46.13 ± 7.74 | 48.69 ± 6.27 | 47.41 ± 6.06 |
| | All | 47.28 ± 7.59 | 49.35 ± 5.97 | 48.32 ± 5.99 |
| SB | F | 52.89 ± 6.86 | 52.16 ± 5.93 | 52.53 ± 5.12 |
| | M | 49.75 ± 5.81 | 47.71 ± 6.42 | 48.73 ± 4.48 |
| | All | 51.59 ± 6.59 | 50.32 ± 6.48 | 50.95 ± 5.19 |
| All | F | 48.21 ± 8.68 | 47.38 ± 9.61 | 47.80 ± 8.17 |
| | M | 40.56 ± 11.12 | 41.20 ± 11.30 | 40.88 ± 10.53 |
| | All | 43.80 ± 10.82 | 43.82 ± 11.02 | 43.81 ± 10.18 |

***Online Resource 26*** *Pairwise comparisons of the inward inclination of maxillary teeth angles between morphs, results from the generalized liner mixed models (glmmTMB), when using ln FL for size.*

| Bones | Morph pairs | Estimates | se | lower. CI | upper. CI | t.ratio | p-value |
|---|---|---|---|---|---|---|---|
| Maxilla | LB-PI | 10.250 | 3.330 | 1.690 | 18.817 | 3.083 | 0.011 |
| | PL-SB | -12.440 | 3.140 | -20.530 | -4.344 | -3.956 | < 0.001 |
| | LB-PL | 15.290 | 3.410 | 6.500 | 24.079 | 4.477 | 0.001 |
| | PI-SB | -7.400 | 2.940 | -14.980 | 0.177 | -2.514 | 0.059 |
| | LB-SB | 2.850 | 3.450 | -6.040 | 11.738 | 0.826 | 0.842 |
| | PI-PL | 5.030 | 2.990 | -2.660 | 12.726 | 1.685 | 0.332 |

***Online Resource 27*** *Pairwise comparisons of the inward inclination of maxillary teeth angles between morphs within sex, results from the generalized liner mixed models (glmmTMB), when using ln FL for size.*

| Bones | Sex | Morph pairs | Estimates | se | lower. CI | upper. CI | t.ratio | p-value |
|---|---|---|---|---|---|---|---|---|
| Maxilla | Females | LB-PI | 9.355 | 3.800 | -0.424 | 19.135 | 2.463 | 0.067 |
| | | PL-SB | -10.602 | 2.710 | -17.567 | -3.636 | -3.918 | **< 0.001** |
| | | LB-PL | 9.636 | 4.040 | -0.773 | 20.045 | 2.383 | 0.081 |
| | | PI-SB | -10.322 | 3.850 | -20.242 | -0.401 | -2.678 | **0.038** |
| | | LB-SB | -0.966 | 4.140 | -11.624 | 9.692 | -0.233 | 0.996 |
| | | PI-PL | 0.280 | 3.75 | -9.386 | 9.947 | 0.08 | 1.000 |

| | | | | | | | | |
|---|---|---|---|---|---|---|---|---|
| | Males | LB-PI | 11.153 | 2.440 | 2.304 | 20.002 | 3.242 | **0.007** |
| | | PL-SB | -14.274 | 4.260 | -25.237 | -3.312 | -3.352 | **0.005** |
| | | LB-PL | 20.941 | 4.270 | 9.953 | 31.929 | 4.906 | **< 0.001** |
| | | PI-SB | -4.486 | 4.260 | -15.441 | 6.469 | -1.054 | 0.718 |
| | | LB-SB | 6.667 | 4.910 | -5.961 | 19.295 | 1.359 | 0.526 |
| | | PI-PL | 9.788 | 3.73 | 0.193 | 19.384 | 2.63 | 0.044 |

***Online Resource 28*** *Pairwise comparisons of inward inclination maxillary teeth angles between sexes within morph, results from the generalized liner mixed models (glmmTMB), when using ln FL for size.*

| Bones | Morph | Pairs | Estimates | se | lower. CI | upper. CI | t.ratio | p-value |
|---|---|---|---|---|---|---|---|---|
| Maxilla | PL | F-M | 11.692 | 2.79 | 6.22 | 17.16 | 4.197 | < 0.001 |
| | PI | F-M | 2.184 | 2.71 | -3.13 | 7.5 | 0.806 | 0.420 |
| | LB | F-M | 0.386 | 2.94 | -5.39 | 6.16 | 0.131 | 0.896 |
| | SB | F-M | 8.019 | 3.54 | 1.07 | 14.96 | 2.267 | 0.024 |
| | All | F-M | 5.570 | 1.02 | 3.560 | 7.580 | 5.45 | < 0.001 |

***Online Resource 29*** *The influence of size, morph, sex, side (left or right) and tooth (1st or 5th tooth) on the inward inclination of maxillary teeth. Analysed with generalized liner mixed model (glmmTMB) using ln C-size for size.*

| Terms | $\chi^2$ | df | p-value |
|---|---|---|---|
| Morph Effect | 259.010 | 3 | < 0.001 |
| Sex Effect | 26.874 | 1 | < 0.001 |
| Bone size | 0.003 | 1 | 0.960 |
| Side | 16.375 | 1 | < 0.001 |
| Tooth | 0.001 | 1 | 0.978 |
| Morph x Sex Interaction Effect | 5.990 | 3 | 0.112 |

| | | | |
|---|---|---|---|
| Morph x size Interaction Effect | 10.225 | 3 | 0.017 |
| Sex x size Interaction Effect | 0.264 | 1 | 0.607 |
| Morph x Tooth | 16.416 | 3 | 0.001 |

***Online Resource 30*** *Pairwise comparisons of the inward inclination of maxillary teeth angles between morphs, results from the generalized liner mixed models (glmmTMB), when using ln C-size for size.*

| Bones | Morph pairs | Estimates | se | lower. CI | upper. CI | t.ratio | p-value |
|---|---|---|---|---|---|---|---|
| Maxilla | LB-PI | 12.370 | 3.33 | 3.800 | 20.930 | 3.72 | 0.012 |
| | PL-SB | -11.250 | 2.51 | -17.710 | -4.790 | -4.48 | < 0.001 |
| | LB-PL | 14.340 | 3.07 | 6.430 | 22.240 | 4.67 | < 0.001 |
| | PI-SB | -9.280 | 2.78 | -16.430 | -2.120 | -3.34 | 0.005 |
| | LB-SB | 3.090 | 2.94 | -4.490 | 10.660 | 1.05 | 0.720 |
| | PI-PL | 1.970 | 2.98 | -5.700 | 9.640 | 0.66 | 0.912 |

***Online Resource 31*** *Pairwise comparisons of the inward inclination of maxillary teeth angles between morphs within sex, results from the generalized liner mixed models (glmmTMB), when using ln FL for size.*

| Bones | Sex | Morph pairs | Estimates | se | lower. CI | upper. CI | t.ratio | p-value |
|---|---|---|---|---|---|---|---|---|
| Maxilla | Females | LB-PI | 11.060 | 3.520 | 1.990 | 20.125 | 3.139 | **0.010** |
| | | PL-SB | -9.020 | 2.460 | -15.340 | -2.694 | -3.672 | **0.002** |
| | | LB-PL | 10.050 | 3.640 | 0.690 | 19.419 | 2.764 | **0.030** |
| | | PI-SB | -10.020 | 3.760 | -19.690 | -0.347 | -2.667 | 0.039 |
| | | LB-SB | 1.040 | 3.580 | -8.170 | 10.243 | 0.290 | 0.992 |
| | | PI-PL | -1.000 | 3.83 | -10.860 | 8.856 | -0.26 | 0.994 |
| | Males | LB-PI | 13.680 | 3.740 | 4.060 | 23.292 | 3.661 | **0.002** |
| | | PL-SB | -13.480 | 3.190 | -21.680 | -5.275 | -4.230 | **< 0.001** |
| | | LB-PL | 18.620 | 3.940 | 8.480 | 28.752 | 4.728 | **< 0.001** |
| | | PI-SB | -8.540 | 3.670 | -17.990 | 0.920 | -2.324 | 0.094 |
| | | LB-SB | 5.140 | 4.090 | -5.380 | 15.660 | 1.257 | 0.590 |
| | | PI-PL | 4.980 | 3.60 | -4.330 | 14.209 | 1.37 | 0.517 |

***Online Resource 32*** *Pairwise comparisons of the inward inclination of maxillary teeth angles between sexes within morph, results from the generalized liner mixed models (glmmTMB), when using ln FL for size.*

| Bones | Morph | Pairs | Estimates | se | lower. CI | upper. CI | t.ratio | p-value |
|---|---|---|---|---|---|---|---|---|
| Maxilla | PL | F-M | 9.528 | 2.38 | 4.86116 | 14.19 | 4.008 | < 0.001 |
| | PI | F-M | 3.586 | 2.95 | -2.21117 | 9.38 | 1.214 | 0.225 |
| | LB | F-M | 0.967 | 2.97 | -4.85412 | 6.79 | 0.326 | 0.745 |
| | SB | F-M | 5.057 | 2.58 | -0.00116 | 10.14 | 1.962 | 0.050 |
| | All | F-M | 4.790 | 0.99 | 2.850 | 6.720 | 4.85 | < 0.001 |

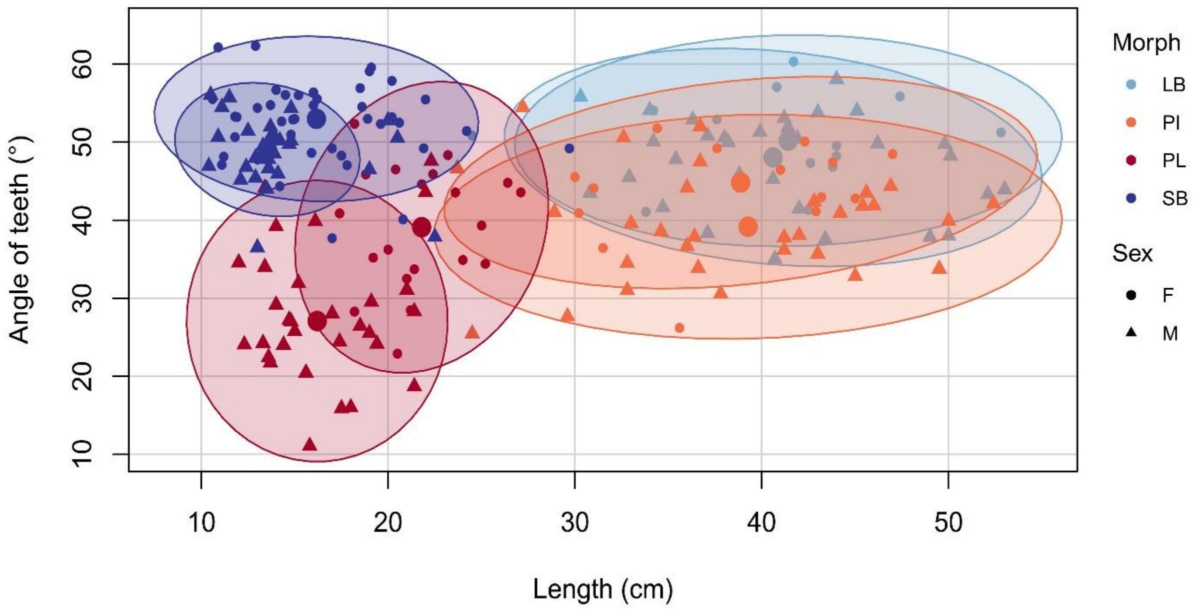


***Online Resource 33*** *Effects of morph and sex on the inward inclination of maxillary teeth in the four morphs. The scatterplot shows average tooth angle (averaged over 1st and 5th tooth on both left and right bone for each individual) (Y-axis) against size (fork length of fish, X-axis). The ellipses and shaded areas are 95% CI for the distributions and the large dots represent the means for the sexes within morphs.*

***Online Resource 34*** *Average number of teeth for six bones and respective standard deviation (SD), calculated for all Arctic charr from the 1997 "replicate" dataset. The numbers represent raw counts.*

| Morph | Sex | Dentary | Maxilla | Premaxilla | Palatine | Vomer | Glossohyal |
|---|---|---|---|---|---|---|---|
| PL | F | 14.80 ± 1.79 | 17.20 ± 1.92 | 7.40 ± 0.55 | 16.00 ± 2.00 | 5.20 ± 1.64 | 3.80 ± 1.10 |
| | M | 16.32 ± 1.94 | 17.38 ± 2.13 | 6.75 ± 1.16 | 15.50 ± 2.53 | 6.10 ± 1.19 | 4.58 ± 1.26 |
| | All | 16.11 ± 1.97 | 17.35 ± 2.07 | 6.84 ± 1.12 | 15.57 ± 2.44 | 5.97 ± 1.28 | 4.47 ± 1.25 |
| PI | F | 16.46 ± 1.98 | 18.23 ± 2.33 | 7.50 ± 1.24 | 16.67 ± 2.96 | 6.58 ± 1.44 | 5.92 ± 1.55 |
| | M | 15.87 ± 2.28 | 19.48 ± 1.92 | 7.41 ± 1.14 | 16.70 ± 2.51 | 6.52 ± 1.44 | 3.81 ± 1.78 |
| | All | 16.08 ± 2.17 | 19.03 ± 2.34 | 7.44 ± 1.16 | 16.69 ± 2.63 | 6.55 ± 1.42 | 4.62 ± 1.97 |
| LB | F | 14.35 ± 1.80 | 19.47 ± 2.53 | 7.24 ± 1.09 | 15.00 ± 2.21 | 6.50 ± 0.97 | 3.94 ± 1.48 |
| | M | 12.73 ± 1.10 | 20.47 ± 2.53 | 6.54 ± 1.05 | 14.29 ± 2.79 | 5.29 ± 0.99 | 4.00 ± 1.31 |
| | All | 13.59 ± 1.70 | 19.94 ± 2.54 | 6.93 ± 1.11 | 14.68 ± 2.47 | 5.93 ± 1.14 | 3.97 ± 1.38 |
| SB | F | 13.00 ± 1.87 | 16.88 ± 2.15 | 7.46 ± 1.22 | 14.00 ± 1.86 | 5.91 ± 1.38 | 4.80 ± 1.68 |
| | M | 12.43 ± 0.53 | 18.00 ± 3.51 | 7.00 ± 1.41 | 13.00 ± 3.03 | 5.43 ± 1.40 | 3.37 ± 1.40 |
| | All | 12.88 ± 1.68 | 17.13 ± 2.50 | 7.38 ± 1.24 | 13.79 ± 2.13 | 5.80 ± 1.37 | 4.53 ± 1.68 |
| All | F | 14.28 ± 2.26 | 17.95 ± 2.47 | 7.40 ± 1.12 | 15.04 ± 2.42 | 6.16 ± 1.35 | 4.73 ± 1.70 |
| | M | 15.12 ± 2.42 | 18.72 ± 6.68 | 6.93 ± 1.18 | 15.44 ± 2.79 | 6.00 ± 1.31 | 4.15 ± 1.47 |
| | All | 14.75 ± 2.38 | 18.38 ± 2.61 | 7.14 ± 1.17 | 15.27 ± 2.63 | 6.07 ± 1.32 | 4.41 ± 1.60 |

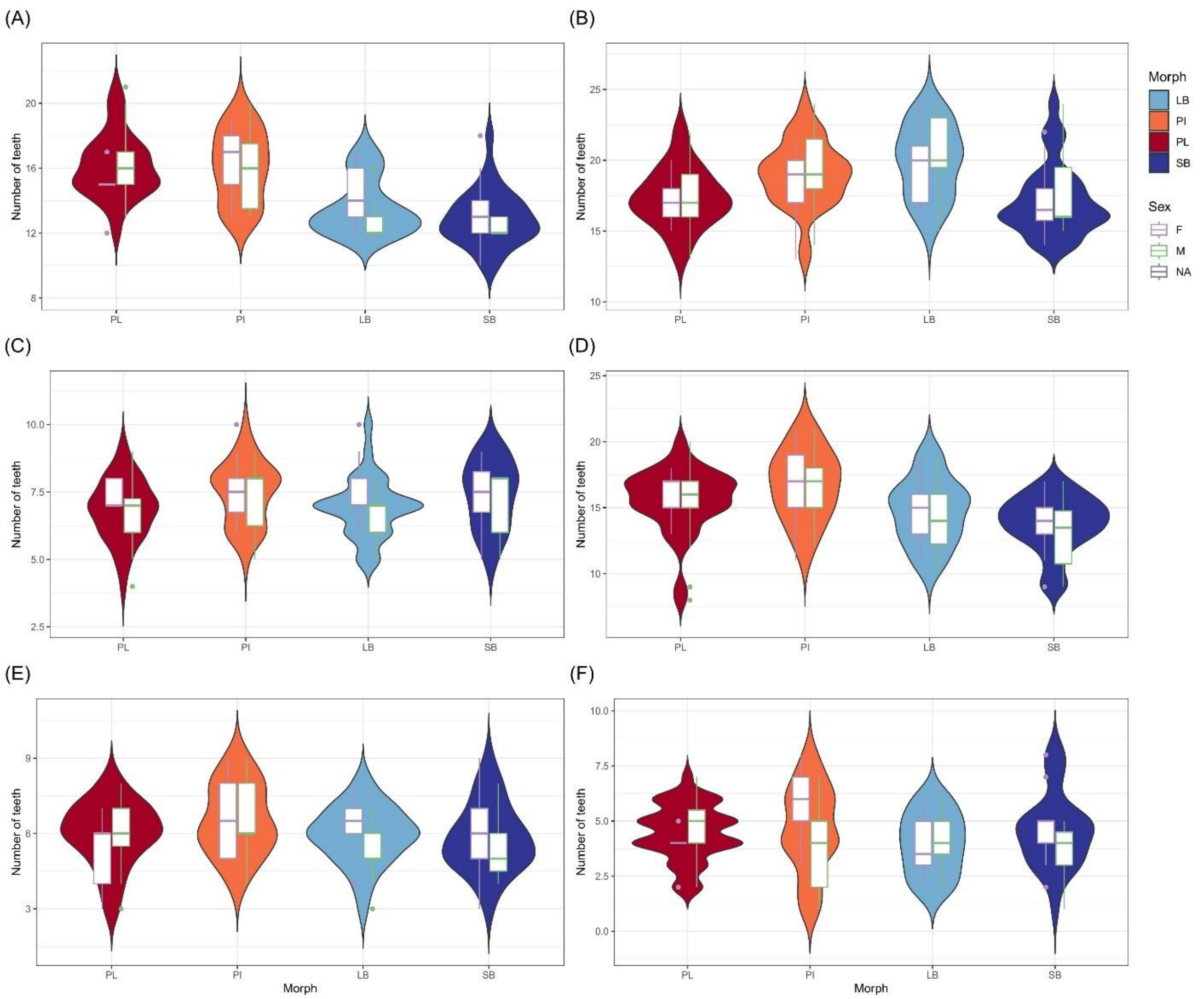


***Online Resource 35*** *Distribution of tooth number in six bones by morph, based on the 1997 "replicate" dataset. Also shown are sex specific distributions as boxplots. The Y-axis indicates the number of teeth. The bones are: dentary (A), maxilla (B), premaxilla (C), palatine (D), vomer (E) and glossohyal bone (F). Morphs are coloured: SB: dark blue, LB: light blue, PI: orange, PL: red (see also inset). Superimposed are boxplots for the sexes (female: purple, males: green).*

***Online Resource 36*** *The influence of size, morph and sex on tooth number in six jaw bones, from the 1997 "replicate" dataset. Analysed with generalized liner mixed models (glmmTMB) using ln FL for size. P-values below the Bonferroni corrected threshold (α = 0.008) are shown in bold.*

| Terms | | Dentary | Maxilla | Palatine | Premaxilla | Vomer | Glossohyal |
|---|---|---|---|---|---|---|---|
| Morph Effect | $\chi^2$ | 75.099 | 10.486 | 20.623 | 5.340 | 9.020 | 9.407 |
| | df | 3 | 3 | 3 | 3 | 3 | 3 |
| | p-value | **< 0.001** | 0.015 | **< 0.001** | 0.149 | 0.030 | 0.024 |
| Sex Effect | $\chi^2$ | 1.793 | 8.814 | 0.624 | 3.415 | 1.232 | 7.021 |
| | df | 1 | 1 | 1 | 1 | 1 | 1 |
| | p-value | 0.181 | **0.003** | 0.430 | 0.430 | 0.267 | 0.008 |
| FL Effect (ln FL) | $\chi^2$ | 0.583 | 1.903 | 0.555 | 0.717 | 8.904 | 7.318 |
| | df | 1 | 1 | 1 | 1 | 1 | 1 |
| | p-value | 0.445 | 0.168 | 0.456 | 0.397 | **0.003** | **0.007** |
| Morph x Sex Interaction Effect | $\chi^2$ | 12.140 | 3.873 | 2.448 | 1.271 | 10.281 | 13.316 |
| | df | 3 | 3 | 3 | 3 | 3 | 3 |
| | p-value | **0.007** | 0.276 | 0.485 | 0.736 | 0.016 | **0.004** |
| Morph x ln FL Interaction Effect | $\chi^2$ | 6.629 | 24.441 | 16.812 | 3.746 | 6.116 | 2.954 |
| | df | 3 | 3 | 3 | 3 | 3 | 3 |
| | p-value | 0.085 | **< 0.001** | **< 0.001** | 0.290 | 0.106 | 0.399 |
| Sex x ln FL Interaction Effect | $\chi^2$ | 2.846 | 2.956 | 2.596 | 0.044 | 0.856 | 1.342 |
| | df | 1 | 1 | 1 | 1 | 1 | 1 |
| | p-value | 0.092 | 0.086 | 0.107 | 0.834 | 0.355 | 0.247 |

P value, Bonferroni corrected threshold, α = 0.008

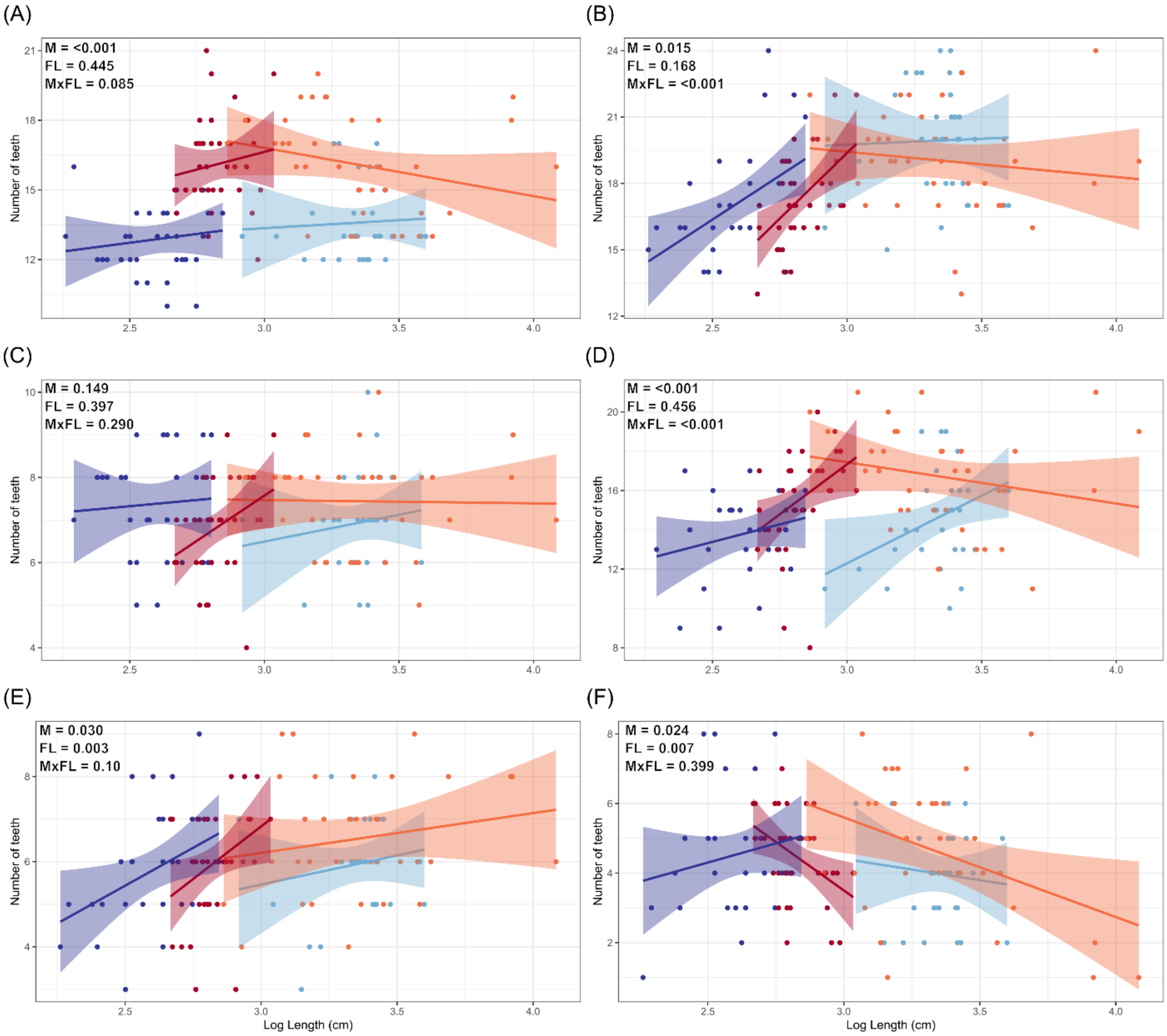


***Online Resource 37*** *Influence of size and morph on number of teeth from the 1997 "replicate" dataset. The scatterplots show individual tooth numbers (Y-axis) against size (ln FL in cm, X-axis) for the four morphs and six bones; A) dentary, B) maxilla, C) premaxilla, D) palatine, E) vomer and F) glossohyal. Each plot shows p-values from glmmTMB for the effect of morph (M), body size (FL) and morph by body size (MxFL) on tooth number.*

***Online Resource 38*** *Average angle of maxilla teeth (± standard deviation) for each morph, also by sex from the 1997 "replicate" dataset.*

| Morph | Sex | Angel (°) |
|---|---|---|
| PL | F | 7.53 ± 8.12 |
| | M | 8.94 ± 9.50 |
| | All | 8.74 ± 9.23 |
| PI | F | 39.22 ± 16.32 |
| | M | 40.07 ± 11.17 |
| | All | 39.77 ± 13.04 |
| LB | F | 51.78 ± 11.63 |
| | M | 52.81 ± 8.39 |
| | All | 52.26 ± 10.10 |
| SB | F | 64.81 ± 7.31 |
| | M | 51.76 ± 8.16 |
| | All | 62.05 ± 9.14 |
| All | F | 51.03 ± 19.66 |
| | M | 30.96 ± 21.28 |
| | All | 39.90 ± 22.81 |

***Online Resource 39*** *The influence of size, morph and sex on the inward inclination of maxillary teeth, from the 1997 "replicate" dataset. Analysed with generalized liner mixed models (glmmTMB) using ln FL for size.*

| Terms | $\chi^2$ | df | p-value |
|---|---|---|---|
| Morph Effect | 370.8672 | 3 | < 0.001 |
| Sex Effect | 1.2712 | 1 | 0.259448 |
| FL Effect (ln FL) | 2.8478 | 1 | 0.091501 |
| Morph x Sex Interaction Effect | 13.0954 | 3 | 0.00435 |
| Morph x ln FL Interaction Effect | 7.6593 | 3 | 0.053604 |
| Sex x ln FL Interaction Effect | 5.3767 | 1 | 0.020407 |

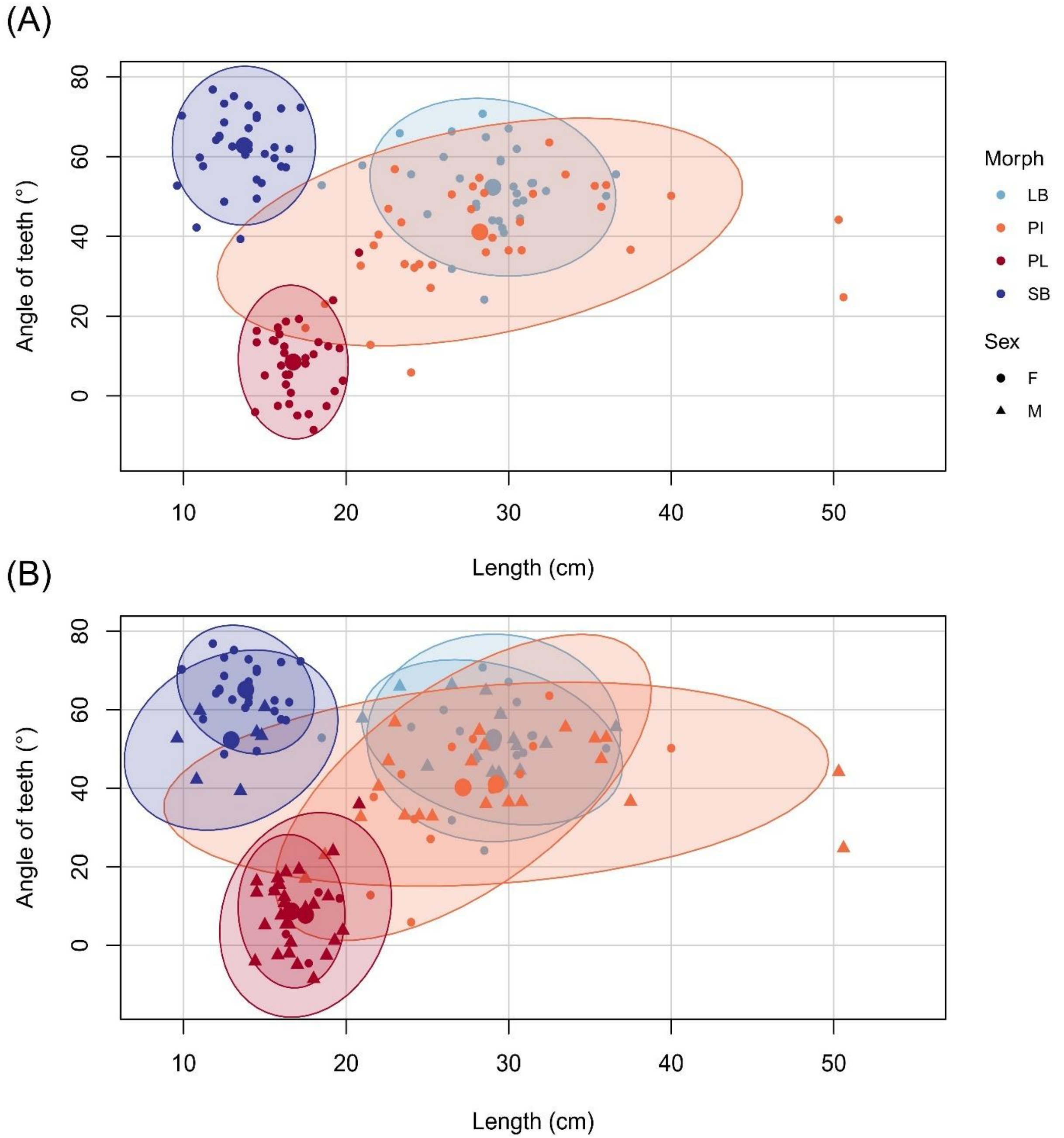


***Online Resource 40*** *Maxilla tooth angle from the 1997 "replicate" dataset varies by morphs (A) and sex by morph (B). The scatterplots show average teeth angle for 1st and 5th tooth on both left and right bone (Y-axis) against size (Fork length of fish, X-axis). Maxilla teeth angle varies by morphs (A) and sex by morph (B). The ellipses and shaded areas are 95% CI for the distributions and the large dots represents the mean for each group. Two datasets were analysed, 240 fish from Jonsdottir 2024, combined data for tooth 1 and 5, with medial/ventral method (see figure 5 and Online Resource 33) and 138 fish from Ingimarsson 2001, measured on tooth 2-5 with fronto-lateral method.*

***Online Resource 41*** *Test of integration between bone shape and tooth number. Traits were compared using partial least squares (PLS), within and among bones.*

| Comparison | Shape | Teeth | r-PLS | z | p-value |
|---|---|---|---|---|---|
| Dentary shape to teeth number | Dentary | Dentary | 0.717 | 5.7832 | 0.001 |
| | Dentary | Maxilla | 0.421 | 4.1109 | 0.001 |
| | Dentary | Premaxilla | 0.113 | -0.8237* | 0.784 |
| | Dentary | Palatine | 0.472 | 4.58 | 0.001 |
| | Dentary | Vomer | 0.419 | 4.0712 | 0.001 |
| | Dentary | Glossohyal | 0.394 | 3.906 | 0.001 |
| | Dentary | Angle | 0.498 | 4.4134 | 0.001 |
| Maxilla shape to teeth number | Maxilla | Dentary | 0.662 | 7.9804*** | 0.001 |
| | Maxilla | Maxilla | 0.486 | 6.6816 | 0.001 |
| | Maxilla | Premaxilla | 0.17 | 0.6525** | 0.253 |
| | Maxilla | Palatine | 0.402 | 4.6623 | 0.001 |
| | Maxilla | Vomer | 0.42 | 5.9191 | 0.001 |
| | Maxilla | Glossohyal | 0.381 | 4.7939 | 0.001 |
| | Maxilla | Angle | 0.448 | 5.7379 | 0.001 |
| Premaxilla shape to teeth number | Premaxilla | Dentary | 0.534 | 7.6948 | 0.001 |
| | Premaxilla | Maxilla | 0.451 | 5.691 | 0.001 |
| | Premaxilla | Premaxilla | 0.203 | 0.5998**** | 0.273 |
| | Premaxilla | Palatine | 0.389 | 4.3004 | 0.001 |
| | Premaxilla | Vomer | 0.411 | 5.0297 | 0.001 |
| | Premaxilla | Glossohyal | 0.395 | 4.6146 | 0.001 |
| | Premaxilla | Angle | 0.521 | 6.1204 | 0.001 |

*Effect size of dentary-premaxilla was significantly lower than all other comparisons.

**Effect size of maxilla-premaxilla was significantly lower than all other comparisons, except maxilla - vomer and maxilla - glossohyal.

***Effect size of maxilla-dentary was higher than maxilla-vomer (p= 0.002)

****Effect size of premaxilla-premaxilla was significantly lower than premaxilla – dentary, premaxilla – maxilla and premaxilla – angle.

***Online Resource 42*** *Test of covariation between bone shape and tooth traits (numbers and angle) by morph. Traits were compared using partial least squares (PLS), within and among (All) morphs. Corresponding pairs of tooth counts and shape are underlined.*

| Covariation | Bone shape | Tooth counts | r-PLS | p-value | | | | |
|---|---|---|---|---|---|---|---|---|
| | | | | PL | PI | LB | SB | All morphs |
| Dentary shape to tooth traits | <u>Dentary</u> | <u>Dentary</u> | 0.418 - 0.689 | **0.001** | 0.503 | 0.082 | **0.001** | **0.001** |
| | Dentary | Maxilla | 0.327 – 0.476 | 0.462 | 0.246 | 0.572 | 0.049 | **0.001** |
| | Dentary | Premaxilla | 0.253 – 0.422 | 0.33 | 0.528 | 0.559 | 0.729 | 0.784 |

| | | | | | | | | |
|---|---|---|---|---|---|---|---|---|
| | Dentary | Palatine | 0.299 – 0.591 | **0.001** | 0.656 | 0.462 | 0.463 | **0.001** |
| | Dentary | Vomer | 0.319 – 0.378 | 0.433 | 0.954 | 0.554 | 0.180 | **0.001** |
| | Dentary | Glossohyal | 0.306 – 0.489 | 0.272 | 0.186 | 0.325 | 0.383 | **0.001** |
| | Dentary | Angle | 0.218 – 0.460 | 0.176 | 0.434 | 0.310 | 0.948 | **0.001** |
| Maxilla shape to tooth traits | Maxilla | Dentary | 0.383 – 0.622 | **0.001** | **0.005** | 0.137 | 0.015 | **0.001** |
| | Maxilla | Maxilla | 0.407 – 0.545 | **0.002** | 0.107 | **0.004** | 0.013 | **0.001** |
| | Maxilla | Premaxilla | 0.285 – 0.428 | 0.335 | 0.080 | 0.249 | 0.394 | 0.253 |
| | Maxilla | Palatine | 0.335 – 0.421 | 0.201 | 0.098 | 0.222 | 0.159 | **0.001** |
| | Maxilla | Vomer | 0.286 – 0.453 | 0.082 | 0.377 | 0.521 | 0.422 | **0.001** |
| | Maxilla | Glossohyal | 0.315 – 0.499 | 0.574 | 0.422 | 0.015 | 0.221 | **0.001** |
| | Maxilla | Angle | 0.267 – 0.497 | 0.024 | 0.686 | 0.325 | 0.608 | **0.001** |
| Premaxilla shape to tooth traits | Premaxilla | Dentary | 0.365 – 0.578 | 0.855 | 0.008 | 0.729 | 0.136 | **0.001** |
| | Premaxilla | Maxilla | 0.374 – 0.531 | 0.506 | 0.051 | 0.853 | **0.003** | **0.001** |
| | Premaxilla | Premaxilla | 0.358 – 0.490 | 0.049 | 0.710 | 0.335 | 0.096 | 0.273 |
| | Premaxilla | Palatine | 0.365 – 0.476 | 0.270 | 0.157 | 0.249 | 0.460 | **0.001** |
| | Premaxilla | Vomer | 0.399 – 0.420 | 0.392 | 0.520 | 0.563 | 0.238 | **0.001** |
| | Premaxilla | Glossohyal | 0.309 – 0.599 | 0.830 | **0.001** | 0.400 | 0.133 | **0.001** |
| | Premaxilla | Angle | 0.365 – 0.552 | 0.026 | 0.499 | 0.646 | 0.595 | **0.001** |

P-value, Bonferroni corrected threshold, α = 0.007
